# The Maintenance of Attention Over Time Influences the Dynamics of EEG Microstates

**DOI:** 10.64898/2026.04.02.716150

**Authors:** Anthony P. Zanesco, Abigail M. Gross, Delanie J. Spivey, Braxton M. Stevenson, Lexi F. Horn, Sophia R. Zanelli

## Abstract

Human attention is inherently transient and limited in span to only a few moments without lapsing. The intrinsic dynamics of large-scale neurocognitive networks are thought to contribute to these lapses and result in the unavoidable fluctuations in attention that constrain its span. However, it remains unclear how the millisecond temporal dynamics of specific electrophysiological brain states contribute to the endogenous maintenance of attention or the onset of attentional lapses. In the present study, we investigated whether the strength and millisecond dynamics of brain electric microstates differentiate states of focus from inattention and contribute to the endogenous maintenance of attention over short and long timescales. We recorded 128-channel EEG while participants maintained their attention during the wait time delay of trials in the Sustained Attention to Cue Task (SACT) and segmented the EEG into a categorized time series of microstates based on data-driven clustering of topographic voltage patterns. The findings revealed that the prevalence and rate of occurrence of microstates C and E in the wait time delay of trials differentiated trials in which the target stimulus was correctly detected from incorrectly detected. These same microstates were also implicated in the maintenance of attention over short and long timescales, with their time-varying dynamics changing systematically during the wait time delay of trials and over the course of the task session. Together, these findings demonstrate the sensitivity of microstates to variation in attentional states and suggest that the millisecond dynamics of these brain states contribute to the maintenance of attention over time.

---

Attention cannot be maintained indefinitely and is limited in span to only a few moments without lapsing or being drawn away from our present goals by distractions or the intrusion of task-unrelated thoughts. Readiness quickly deteriorates and stimulus detection accuracy begins to be impaired within seconds after attention is directed and endogenously maintained on a location in space before reaching its lower limit tens of seconds later (Tsukahara et al., 2023; Zanesco et al., 2026b). Attempts to sustain attention over longer durations reliably lead to worsening performance as time progresses over the course of task sessions (e.g., See et al., 1995; Schwartzman et al., 2025; Zanesco et al., 2024) alongside increasing rates of occurrence of internally generated task-unrelated thoughts (Zanesco et al., 2025). The neurocognitive processes contributing to the maintenance of attention and its lapses are thought to be dynamic and reliant on coordinated neural circuits that demonstrate intrinsic time-varying fluctuations in their activity. These network dynamics may result in natural, unavoidable fluctuations in attention that contribute to momentary behavioral errors and constrain our attention span to short moments.

Converging evidence from electrophysiological and functional neuroimaging studies demonstrates that the brain circuits supporting goal-directed attention are not static in their levels of activity but instead predominate temporarily before transitioning to other active brain network configurations. Fluctuations in activity of a set of distributed attention-related frontoparietal regions covary with the occurrence of momentary attentional lapses, and, importantly, coincide with weaker task-related suppression of the default mode network (Weissman et al., 2006). Infraslow fluctuations in the activity of the dorsal attention and frontoparietal control networks in tasks requiring external attention are also inversely correlated with activity in the default mode network, and stronger decoupling between these networks is associated with better sustained attention (Abbas et al., 2019; Seeburger et al., 2024; Watters et al., 2025). Indeed, the default mode network is thought to exhibit an antagonistic, anti-correlated relationship with brain systems supporting attention oriented towards sensory stimuli; its activity waxes and wanes as attention and sensory systems vary in priority (Fox et al., 2005; Menon, 2023; Raichle, 2015; Smallwood et al., 2021). These patterns highlight the inherently dynamic nature of attention and time-varying competition between the large-scale brain networks mediating its functioning, whose intrinsic variability enables flexibility in how attention is deployed but may also impose limits on its span.

The default mode network is thought to play an important role in supporting internally oriented cognition, such as when attention shifts away from the external sensory environment to internally generated task-unrelated thoughts during episodes of mind wandering (Andrews-Hanna et al., 2014; Christoff et al., 2009, 2016; Fox et al., 2015; Kucyi et al., 2023; Smallwood et al., 2021). Principal network nodes are positioned at the extreme end of a cortical functional gradient separating its transmodal association regions from sensory-motor regions (Margulies et al., 2016; Huntenburg et al., 2018). Timescales of cortical electrical rhythms recapitulate this gradient with the intracranial peak spectral frequencies in transmodal regions demonstrating slower frequency profiles than sensory-motor regions (Royer et al., 2026). These perspectives imply that one of the brain’s organizing principles is structured to insulate higher-order and internally oriented cognitive processes from sensory input. This principle is also consistent with research demonstrating that perceptual processing of external stimuli is attenuated when attention turns inward towards task-unrelated thoughts during the occurrence of mind wandering (Baird et al., 2014; Denkova et al., 2018; Franklin et al., 2013; Kam et al., 2011, 2022; Smallwood, 2013; Smallwood et al., 2008).

Electrophysiological evidence also confirms that attention, perception, and the brain systems supporting these processes fluctuate at timescales orders of magnitude faster than can be identified based on the slow hemodynamics of the brain (Fiebelkorn et al., 2013, 2018; Helfrich et al., 2018; VanRullen, 2016). For example, electroencephalographic (EEG) recordings suggest that ongoing oscillatory rhythms in scalp-recorded electrical activity shift with attentional states and the occurrence of mind wandering. Greater pre-stimulus power in the alpha (8–14 Hz) frequency band precedes failures to detect visual stimuli in tasks requiring perceptual decisions (e.g., Linkenkaer-Hansen et al., 2004; Michail et al., 2022; Samaha et al., 2020; van Dijk et al., 2008) and greater alpha power tends to co-occur with the moments of mind wandering compared to on-task states (see Kam et al., 2022, for review). Elevated alpha power, particularly at posterior electrode sites, is increasingly viewed as a marker of reduced cortical excitability that limits the processing and conscious perception of sensory stimuli (Samaha et al., 2020). These reductions in cortical excitability are thought to produce non-specific changes in stimulus detection by shifting detection criterion rather than sensitivity for distinguishing between different stimuli because the processing of all sensory inputs is affected in those moments (Samaha et al., 2022).

The millisecond dynamics of momentarily predominating global brain states are also evident in the scalp-recorded EEG and these microstates have been shown to differentiate states of inattention from focus in a continuous performance task (Zanesco et al., 2021). Microstates represent brief electrophysiological events identifiable from distinct topographic patterns of voltage across the scalp that originate from large amplitude co-fluctuations in the neuronal activity of coordinated brain circuits (Michel & Koenig, 2018). Each topographically unique microstate tends to persist momentarily for roughly 30–120 msec before transitioning to other configurations (Zanesco, 2024). The presence of different microstates can differentiate the activity of distinct neural populations because topographic variations in their configuration must reflect differences in the spatial distribution of their generators. The topographic configurations of microstates are highly replicable with roughly four to seven microstates accounting for the majority of topographic variance in spontaneous EEG (Koenig et al., 2024). This generalizability suggests a neural repertoire of brain states originating from a common brain network architecture in humans.

The prevalence of two specific microstates occurring in the pre-stimulus period of task trials, each distinguishing the activity of unique sets of brain generators, were found to be associated with mind wandering and task focus (Zanesco et al., 2021). Specifically, the prevalence, strength, and frequency of occurrence of microstate configuration C was greater in the moments preceding experience sampling probe-caught reports of being unfocused and “off-task,” whereas microstate configuration E was greater preceding reports of being “on-task.” These patterns are reminiscent of the supposed antagonistic interplay between brain systems supporting goal-directed attention and the default mode network, suggesting the possibility that microstates may originate from the coordinated electrophysiological activity of large-scale functional brain networks. In support of this view, tentative evidence suggests that the distributed sources underlying microstates overlap with key regions of fMRI-defined functional networks, including the default mode network (Bagdasarov et al., 2022; Brechet et al., 2019; Britz et al., 2010; Custo et al., 2017; Ngo et al., 2026; Valt et al., 2024; Tarailis et al., 2025; Zanesco et al., 2026a).

Nevertheless, the brain generators and functional relevance of these momentarily predominating brain states are still uncertain because studies have primarily investigated microstates during resting-state EEG recordings rather than in cognitive-behavioral task contexts.

The brain generators of microstates contribute substantial amounts of power within broadband oscillatory frequencies due to periodic and aperiodic fluctuations in their activity. Indeed, the average duration and occurrence of microstates are largely explained by periodicities in the dominant alpha frequency band (Lehmann et al., 1987; Milz et al., 2017; von Wegner et al., 2017, 2021; Wiemers et al., 2024). Furthermore, their dynamics resemble the behavior of standing waves (Michel & Koenig, 2018; von Wegner et al., 2021), as the same relative topographic configuration can often be seen to oscillate in polarity for several cycles before a different microstate takes its place. The connectivity repertoires that mediate sequential transitions between different microstate configurations can themselves be explained by propagation of traveling waves in the alpha band (Li et al., 2025). While the brain generators of microstates contribute prominently to alpha band power on the scalp, it is also clear that microstates are not reducible to specific band-limited oscillations and instead demonstrate complex aperiodic and time-varying dynamics between bouts of rhythmicity (Michel & Koenig, 2018).

In the present study, we employed a modified version of the Sustained Attention to Cue Task (SACT; Zanesco et al., 2026b) to examine the activity and dynamics of EEG microstates and their functional relevance for sustained attention and the occurrence of mind wandering. The SACT requires attention to be endogenously maintained at a cued spatial location to detect a briefly presented target stimulus amidst an array of distractors. The time that attention must be maintained is parametrically varied on task trials so that the demand placed on sustained attention during this wait time delay is distinct from change occurring over the entire task session. Data-driven clustering identified the distinct topographic microstates occurring during the wait time delay of trials from high-density EEG recorded in the task. The activation strength of microstates and their millisecond temporal dynamics were examined to evaluate their association with attentional lapses and change during the wait time delay or over the course of the task session. This investigation aims to investigate how the millisecond dynamics of microstates contribute to intrinsic constraints on the endogenous maintenance of attention.

The SACT was developed to measure sustained attention as part of a comprehensive toolbox of attention control tasks (Draheim et al., 2021, 2024; Tsukahara et al., 2023) and is an analog of the Psychomotor Vigilance Task (Dinges & Powell, 1985) that relies on accuracy rather than response time as the primary outcome measure of behavioral performance. Prior studies have found that target detection accuracy substantially worsens on trials with longer wait time delays (Tsukahara et al., 2023; Zanesco et al., 2026b). Experience sampling mind wandering probes also caught individuals experiencing worse self-reported focus and more task-unrelated thought on trials with longer wait times or later in the task session (Zanesco et al., 2026b). Unlike other continuous performance tasks used to investigate sustained attention, the maintenance of attention during the wait time delay of the SACT relies primarily on endogenous attentional control because external stimuli do not interrupt the delay period. This provides an ideal opportunity to experimentally isolate the capacity limitations of endogenous attention and its associated brain dynamics without interference from stimulus-related brain activity.

In line with prior research identifying specific microstates that distinguish states of inattention from focus (i.e., Zanesco et al., 2021), we predicted that the activity and dynamics of two distinct microstates, configurations C and E, would be associated with target detection errors and reports of mind wandering in the SACT. We expected microstate C to be more prevalent and occur more frequently when individuals were inattentive during the wait time delay of trials. This prediction aligns with prior research demonstrating that microstate C is more present in the pre-stimulus periods of trials preceding episodes of mind wandering (Zanesco et al., 2021) and studies suggesting its generators within subsystems of the default mode network (e.g., Custo et al., 2017). Accordingly, we expected microstate C to grow in prevalence over the task session, in parallel with increasing rates of mind wandering over time across a range of cognitive-behavioral tasks (Zanesco et al., 2025) and in the SACT (Zanesco et al., 2026b). In contrast, we expected microstate E to be more prevalent and occur more frequently when individuals maintained their focus over the wait time delay. As prior neuroimaging studies have shown increased activity in distributed frontoparietal regions on trials when attention must be maintained over longer inter-target intervals (Breckel et al., 2011; Langner & Eickhoff, 2013), we predicted that microstate E would be more prevalent and occur more frequently over longer wait time delays.

We also had several secondary exploratory aims for this investigation. First, we sought to investigate the microstate-correlates of accumulating attentional demands by evaluating the influence of previous trial wait times on current trial microstates. Microstates may provide insight into the mechanisms through which target detection accuracy was worse in the SACT when the current trial was preceded by trials with longer wait time delays (Zanesco et al., 2026b). Second, we quantified the distributed neural generators of microstates using EEG source localization to evaluate the degree to which their generators overlap with large-scale brain networks and attention-related brain regions previously characterized using functional neuroimaging. Third, we evaluated whether the activity and dynamics of microstates in the wait time delay of trials can account for associations between pre-stimulus alpha power and states of inattention. Accordingly, the brain generators of specific microstates, such as microstate configuration C, may be responsible for the presence of greater alpha power in pre-stimulus periods preceding stimulus detection errors and the occurrence of mind wandering. This would contribute to debates (i.e., Samaha et al., 2020) regarding the mechanisms by which alpha power influences target detection accuracy by identifying the momentary brain states and their underlying generators that contribute to task-relevant differences in alpha power.

## Methods

### Participants

Fifty-one individuals (M age = 19.59 years, SD = 3.90, age range = 18–45) were recruited from the University of Kentucky undergraduate community to participate in the study. One additional individual was recruited for pilot testing of study procedures, but their data is not included in the study. Participants were predominately female (n = 37), non-Hispanic (n = 45), and most reported a White racial identity (n = 40), whereas the remaining participants reported an Asian or Asian American identity (n = 2), a Black or African American identity (n = 4), or another unspecified identity (n = 5). Participants verbally confirmed that they had normal or corrected-to-normal vision, and felt alert, rested, and received an adequate amount of sleep the prior evening (M = 7.83 hours of sleep, SD = 1.10). No participant reported a history of medical or neurological diagnoses that would prevent the completion of any activities in the session. Forty-nine individuals comprised the final sample size after participant exclusions (see Procedure). The Institutional Review Board of the University of Kentucky approved the study. Participants provided written informed consent and received course research credit for participation.

### Procedure

Participants completed a series of questionnaires and a modified version of the Sustained Attention to Cue Task (SACT; Draheim et al., 2021, 2024). Participants sat alone in a quiet, illuminated room and completed the questionnaires and the task on a Windows 11 computer with a 24-inch monitor. The task was administered using E-Prime 3.0 (Psychology Software Tools, 2016). Compliance during the session was monitored through a half-silvered mirror by an experimenter in an adjacent room. Two participants were subsequently dismissed from the study and excluded from all analyses because they were observed to be falling asleep during completion of the SACT.

#### Sustained Attention to Cue Task

A modified version of the SACT was administered to participants as part of the experimental session. Attention must be maintained in the SACT over a variable wait time interval to correctly detect target stimuli at spatially cued locations. On each trial, participants are cued to attend to a spatial location on the blank monitor screen (grey background) and maintain their attention at the cued location for a variable amount of time (0 to 40 seconds) until a letter array appeared at the cued location (see Figure 1 for a schematic depicting the sequence and timing of trial events). The cued location could be present within one of four possible quadrants of the display with equal probability in the top-left, top-right, bottom-left, and bottom-right quadrants. Wait times (M = 20 seconds, SD = 13.26) ranged from 0 to 40 seconds. The wait time on each trial was selected randomly from 11 unique wait times (i.e., wait times of 0s, 4s, 8s, 12s, 16s, 20s, 24s, 28s, 32s, 36s, 40s).

**Figure 1.**
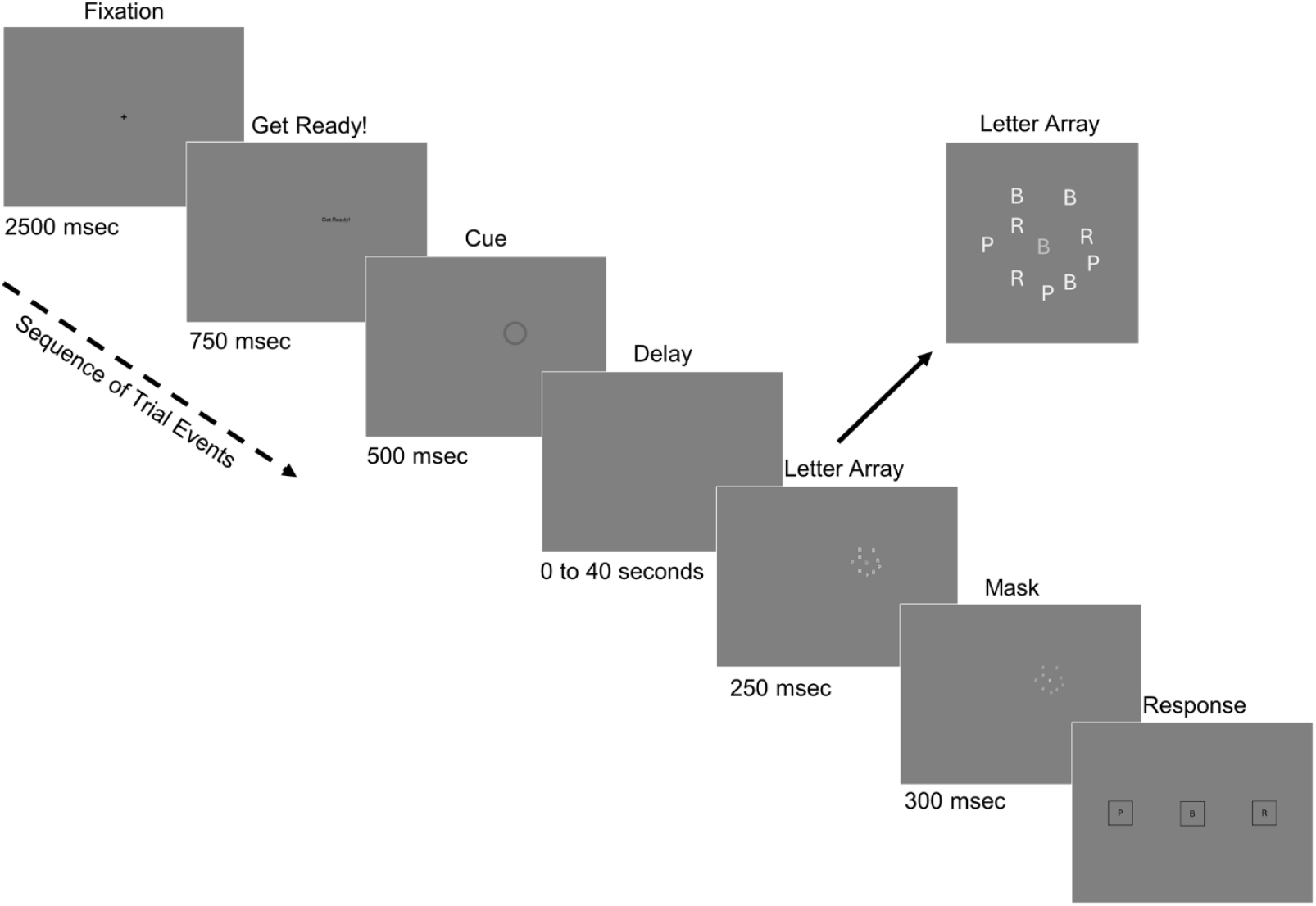
Schematic depicting a trial of the modified Sustained Attention to Cue Task (SACT). Participants are cued to attend to a spatial location on the blank monitor screen (grey background) and maintain their attention at the location for a variable wait time delay (0 to 40 seconds) until a letter array appears at the cued location. Participants then select what they thought was the correct target letter at the center of the letter array by choosing the corresponding option on the response screen. Experience sampling mind wandering probes replaced the presentation of the letter array on a subset of trials.

An array of letters appeared for 250ms at the cued location at the end of the wait time interval. Participants had to identify the target letter at the center of the array presented in dark gray font. Nine non-target distractor letters surrounded the target in silver font. The target was either a “B,” “P,” or “R” and the nine non-target letters were equal numbers of each of those letters. The target letter occurred at the center of the cued location. Non-target letters surrounded the target and were semi-randomly dispersed within a 96 x 96-pixel square. Stimuli had a minimum distance of 24 pixels. After the letter array, a mask (the hash symbol “#”) replaced each letter for 300ms followed by a response screen with “B,” “P,” and “R” response options. Participants selected what they thought was the correct target letter by clicking the corresponding option on screen using a mouse. The response screen was displayed until a response was provided. A blank monitor screen was displayed for 500 msec at the end of each trial.

Mind wandering experience sampling probes were presented on a subset of trials. At the end of the wait time interval on these trials, text appeared that instructed participants to rate a series of four successive statements describing their attentional focus and whether they experienced task-unrelated thought. The rating screen was displayed for each statement until a response was provided. For the first statement, participants rated whether “my attention was focused at the location of the target” by selecting an option ranging from 1 (“not at all focused”) to 5 (“extremely focused”) on the keyboard. For the second statement, participants rated whether “my attention was ready for the target to appear” by selecting an option ranging from 1 (“not at all ready”) to 5 (“extremely ready”). For the third statement, participants rated whether “my attention was focused on my thoughts instead of the task” by selecting an option ranging from 1 (“not at all focused on thoughts”) to 5 (“extremely focused on thoughts”). For the fourth statement, participants rated whether “my thoughts were distracting me from the task” by selecting an option ranging from 1 (“not at all distracting”) to 5 (“extremely distracting”). Probes always occurred on trials with wait times of either 4s, 12s, 20s, or 40s. The letter array never appeared on these trials.

The SACT consisted of five practice trials with feedback about accuracy. Participants were required to get three practice trials correct before continuing. One additional practice trial included a mind wandering probe to familiarize participants with the experience sampling statements. There was no feedback on the remaining task trials. The main task had four blocks of 15 trials for a total of 60 trials. There were 44 total trials with letter arrays and 16 trials with mind wandering probes. Each unique wait time delay was selected only once for each block of task trials for a total of 4 trials per wait time for letter arrays trials and mind wandering probe trials, respectively. There was a self-timed break provided after each block of trials. In total, the task took roughly 32.45 minutes (SD = 1.83) to complete on average (excluding time for instructions and practice).

#### Confirmatory factor analysis of mind wandering ratings

In line with our prior work (Zanesco et al., 2026b), we evaluated whether the four mind wandering ratings were best represented by two distinct facets of mind wandering. Confirmatory factor analysis revealed that two correlated latent factors representing focus and task-unrelated thought were a good fit for the four mind wandering ratings, χ2(2) = 5.361, p = .069, CFI = 0.998, RMSEA = 0.046 [0.000, 0.096]. Coefficient omega calculated at the within-person (level 1) and between-person (level 2) levels suggest high-levels of reliability for factors of focus (level 1 ω = 0.835; level 2 ω = 0.958) and task-unrelated thought (level 1 ω = 0.842; level 2 ω = 0.989). The two factors were more strongly correlated at the within-person (r = −0.732) than the between-person (r = −0.589) level. Accordingly, we calculated scores for focus and task-unrelated thought by averaging the two corresponding mind wandering ratings representing each factor. Focus and task-unrelated thought composite scores were separately analyzed.

### EEG Data Acquisition and Processing

EEG was continuously recorded during the SACT from 128 Ag/AgCl active electrodes using a BioSemi ActiveThree system with a sampling rate of 1024 Hz. Electrodes were located on the scalp according to an equiradial montage. Individual electrode locations were digitized for each participant in three-dimensions using a Polhemus Patriot digitizer (http://www.polhemus.com). Four additional electrodes were placed next to the outer canthi and below the left and right eyes to record horizontal and vertical electrooculograms. EEG were processed offline using the free Cartool software toolbox version 5.06.04 (Brunet et al., 2011). Recordings were downsampled to 256 Hz, bandpass filtered between 1–40 Hz, and average referenced. EEG were visually screened for channels with intermittent connectivity or excessive periods of extreme amplitude, and these channels were interpolated using 3D spline interpolation. Individual electrode locations were also transformed into a standard 128-channel equiradial montage through the same interpolation.

EEG data were then segmented into contiguous 4000 msec epochs collected from the wait time delay of task trials. For example, four contiguous 4000 msec epochs were segmented from the wait time delay of a trial with a 16 second wait time. Trials with wait time delays of 0 seconds were excluded from subsequent processing and analyses of EEG. As such, there were 296 epochs available for analysis for each individual participant (14,479 epochs in total across all participants). Trial event timing and information about the ordering of epochs of each wait time delay was retained after epochs were segmented. Infomax-based Independent Component Analysis (ICA; Jung et al., 2000) was subsequently used to remove non-neural signal contaminants from the EEG, including sources of high-frequency noise, ocular, cardiac, and muscle artifacts. Epochs were separately concatenated for each participant and submitted to ICA. Trial epochs were again separated after ICA. Each epoch was visually inspected for any remaining periods of excessive noise or artifacts, and these periods were marked for exclusion. Finally, the EEG were spatially smoothed to reduce the influence of outlier channels in the voltage topography during topographic clustering and microstate labeling (see Michel & Brunet, 2019).

### Topographic Clustering and Microstate Parameter Estimation

An adapted k-means clustering procedure determined the optimal number of representative topographic clusters of microstates. The aim of clustering was to identify the fewest number of k clusters accounting for the greatest global explained topographic variance (GEV) in the EEG time series (Michel et al., 2009; Murray et al., 2008). A schematic of the clustering procedure is depicted in Figure 2 alongside the results of clustering applied to these data. EEG data during the wait time delay of trials were subsequently discretized into symbolic microstate sequences based on their alignment with the topographic configurations derived from clustering. This enabled the calculation of temporal parameters describing the dynamics of microstates for subsequent statistical analysis. The clustering and categorization of microstates from EEG followed widely employed procedures that have been extensively described (see Zanesco et al., 2020, 2021, 2026a) but the methodological details are nevertheless summarized again in the following sections.

**Figure 2.**
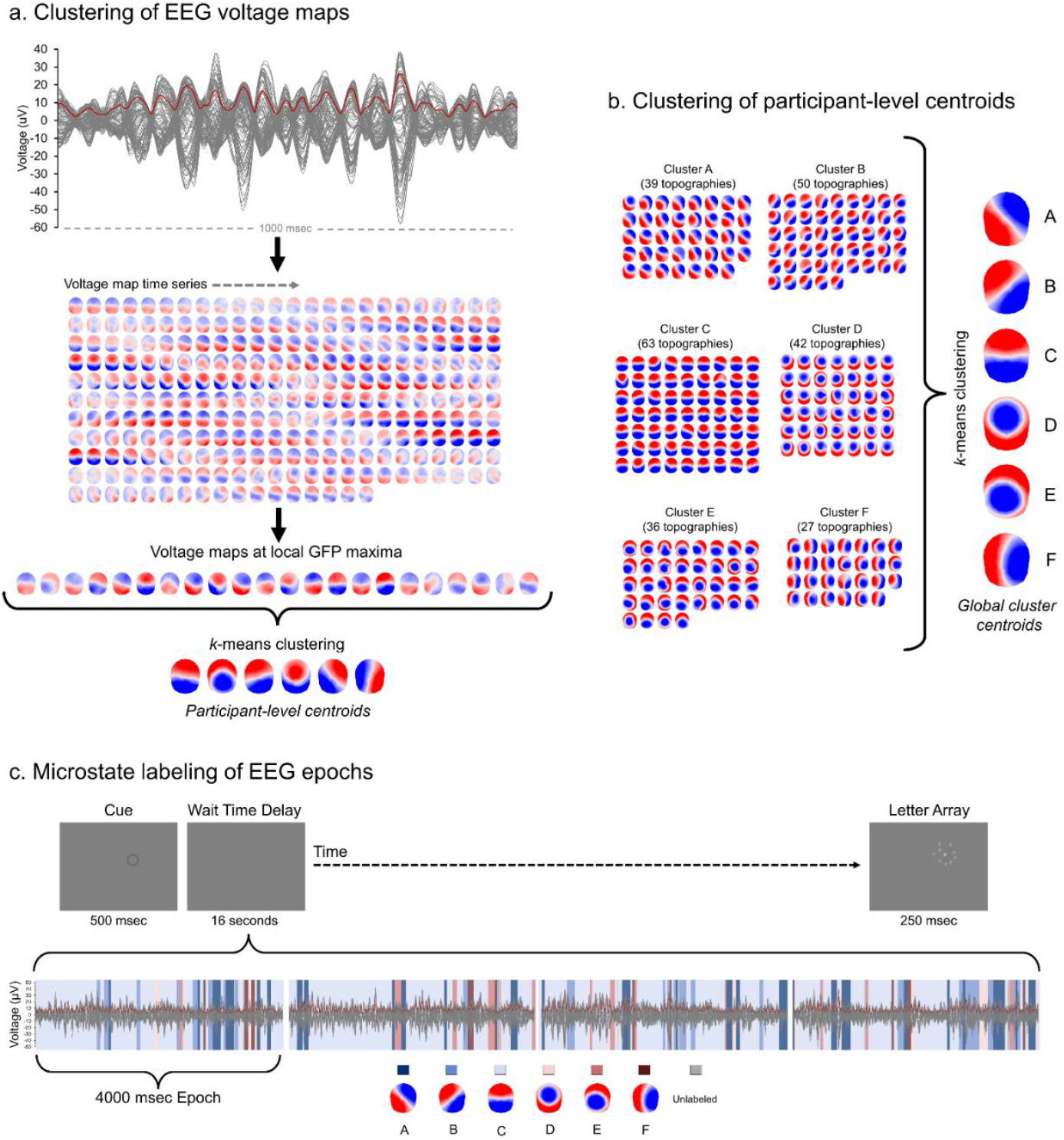
Topographic clustering and microstate labeling of EEG during task epochs. a). Voltage maps at local maxima of the global field power (GFP) are identified from the EEG time series. The GFP time series is shown in red superimposed on a butterfly plot of the 128-channel EEG. The time series of voltage maps are shown below as 2-D isometric projections with nasion upwards. Initial k-means clustering identifies the optimal participant-level topographic clusters. b). The centroids of clusters from participants undergo a second *k*-means clustering to define the global microstate clusters (A through F). Centroids (257 in total) derived from *k*-means clustering of 49 participant recordings are shown grouped by their global cluster membership. c). The microstate time series of epochs during the wait time delay of trials are categorized according to the global microstate configurations. The categorized time series is shown overlaid with the EEG butterfly plot and GFP. Dependent measures of microstates are calculated from the categorized sequence for every epoch and each microstate configuration.

#### Topographic clustering of voltage maps

Topographic voltage maps from sets of EEG epochs at the local maxima in the global field power (GFP) time series were identified for clustering because these moments provide optimal representations of the topography of microstates (Zanesco, 2020). Clustering occurred separately for each individual and included all epochs of trials from that session. GFP is a reference-independent measure of voltage potential (μV) that quantifies the strength of the scalp electric field independent of its topographic configuration (Skrandies, 1990). The adapted k-means clustering procedure assigned voltage maps at the local GFP maxima to k clusters based on their spatial correlation in an iterative process until the GEV converged to a limit. This was repeated for each level of k [1:12] with 100 repetitions per level of k. Maps were only assigned to a cluster if the spatial correlation with the centroid exceeded 0.5; and correlation values were based on the relative topographical configuration but not the polarity of the maps by correcting the sign of the spatial correlation coefficients during cluster assignment (Michel et al., 2009). The optimal number of k clusters was selected using a metacriterion defined by seven independent optimization criteria (see Brechet et al., 2019; Custo et al., 2017). Four to eight topographies (M = 5.25, SD = 0.99) were identified as the optimal number of k clusters among the 49 individuals, explaining 77.35% (SD = 4.98, range = 64.74–90.68) of topographic variance in individuals EEG recordings on average.

After identifying participant-level cluster centroids, we conducted a second stage of k-means clustering to identify the optimal global clusters that best explain the participant-level cluster centroids (totaling 257 topographies) across all 49 participants. Clustering was repeated for each level of k [1:15] with 200 repetitions per k. The optimal number of k global clusters was determined using the optimization metacriterion: six global clusters explained 85.30% of topographic variance (i.e., GEV) among the participant-level cluster centroids. The six clusters were labeled configurations A through F according to their ordering and similarity to the meta-analytic topographic cluster solution reported by Koenig et al. (2024). Each of the six clusters were clearly identified (spatial correlations > 0.92) among the centroids of the meta-analytic clusters (Koenig et al., 2024). Figure 2b depicts the six global topographic configurations alongside the 257 participant-level cluster centroids grouped according to their global cluster membership.

#### Parameterization of the microstate time series

Each time series sample of the multichannel EEG were then categorized according to the six global topographic configurations that most strongly correlated with the GFP normalized voltage map at that moment. This resulted in a symbolic time series of categorical labels representing sequences of microstates for each epoch. EEG samples that correlated poorly with global topographies (spatial correlations < 0.5) were left uncategorized. Polarity was also ignored during categorization by correcting the sign of the spatial correlation coefficient. Finally, temporal smoothing was applied to the symbolic sequence using strength 10 on half window size of 2 (Besag factor k = 10 and b = 2; Pascual-Marqui et al., 1995) and by ignoring assigned microstates that were present for less than 6 consecutive samples (i.e., < 23 msec), then splitting the time points between the preceding and subsequent microstates in the series. 96.20% (SD = 4.73, range = 20.63–100) of time series samples from the 14,479 total epochs were categorized according to the six global microstate topographic configurations after temporal smoothing. The six representative microstate configurations in total explained 60.47% (SD = 6.59, range = 33.97–83.46) of the global explained topographic variance in the EEG time series on average across all 14,479 epochs.

Five dependent measures of microstates were subsequently derived from the symbolic time series of microstate labels for each 4000 msec epoch of EEG: global explained variance (GEV) reflects the percentage of topographic variance in the EEG recording that is explained by a given microstate configuration; mean microstate duration is the average duration (in msec) of contiguous samples categorized according to a specific microstate configuration; mean occurrence rate represents the average times per second (Hz) a given microstate occurs in the EEG time series; number of time samples represents the total time occupied by a given microstate in the EEG time series expressed in time samples; and mean global field power (GFP) is the average of the GFP for all samples categorized according to each microstate. Finally, the probabilities of microstates to transition from one configuration to another in the EEG time series of epochs were calculated for each pair of microstates.

### Source Localization of Microstates

We estimated the neural sources of microstates using a distributed linear inverse solution (LAURA; Grave de Peralta Menendez et al., 2004). Only time series samples with spatial correlations r > .80 with the corresponding cluster centroid were submitted for source localization because these samples best represent each microstate configuration and exclude periods of instability in the voltage topography during momentary transitions between microstates. 52.05% (SD = 8.86) of samples of epochs met this criterion. The 128 electrode locations were co-registered with the Montreal Neurological Institute template brain. The lead field was derived from a 4-shell Local Spherical Model with Anatomical Constraints (LSMAC) head model with 6926 solution points distributed in the grey matter of the MNI template (see Brunet et al., 2011). The average age of the sample was used to estimate skull thickness for the model. Results were optimized with regularization and normalized to reduce biases due to power variability across time series samples (see Michel & Brunet, 2019). The amplitude of the norm of dipoles were obtained as scalar, positive values for solution points. Inverse solutions for all time series samples were averaged according to the categorized microstates at those moments. Solution points were evaluated if they were above the 95th percentile for each microstate configuration of the mean source maps. The results of source localization are provided in Supplementary Materials.

### Microstate Averaged Time-Frequency Transformation

Time-frequency transformation (“wavelet” S-Transform implemented in Cartool; Brown et al., 2010) of the spatially smoothed EEG time series of epochs determined the power in the alpha (8-14 Hz) frequency bands for all electrodes. This enabled estimation of localized variations of alpha power in time within each time series of epochs. Power in this band was subsequently averaged for each epoch according to the microstate configurations categorizing the time samples of the EEG. Power at uncategorized periods of time was ignored. The result was an estimate of power for all electrodes and microstate configurations for each of the 14,479 epochs. Episodes of sustained, large amplitude alpha oscillations were also identified using an approach motivated by methods for identifying oscillatory episodes from wavelet transformed EEG (Hughes et al., 2012; van Vugt et al., 2007). The number of time samples that distinct microstates were present within these oscillatory episodes was quantified and compared between configurations. The results from these analyses are summarized in Supplementary Materials.

### Analysis

#### Behavioral performance and mind wandering

Growth curve models were used to characterize within-task differences in SACT accuracy and probe-caught ratings of mind wandering as a function of the fixed and random effects of trial wait time and time-on-task (i.e., sequential trial number). Growth models were estimated with maximum likelihood in the multi-level mixed effects modeling framework (McNeish & Matta, 2018) using the PROC MIXED procedure in SAS 9.4. Accuracy was aggregated for participants across sets of trials for each unique wait time and values were nested within individuals for analyses of the effects of trial wait time using linear mixed effects models. We also examined trial-level mind wandering ratings as a function of the fixed and random effects of trial wait time and sequential trial number using linear mixed effects models. Model fixed effects describe the average starting point (i.e., intercept) and linear (or polynomial) rate of change (i.e., slope) across trial wait times or sequential trials. Random effects provide estimates of between-person variability and covariance in model parameters. Linear and polynomial fixed effects were included in models when significant. Random slopes for these parameters were included in models if they were successfully estimated.

#### Microstate strength and temporal dynamics

Growth models were used to characterize differences in the five microstate dependent measures derived from EEG epochs as a function of the fixed effects of trial accuracy (correct vs. incorrect trials), sequential epoch number (i.e., time during the wait time delay), and sequential trial number (i.e., time-on-task). Fixed effects of microstate configuration (A through F) and two-way interactions between microstate configuration and other effects were included in models. Fixed effects of trial accuracy (0 = “incorrect” and 1 = “correct”) reflect differences between trials in which individuals correctly detected the target stimulus amid the letter array relative to incorrect trials. Fixed effects of sequential epoch and sequential trial number describe the average starting point (i.e., intercept) and linear rate of change (i.e., slope) across sequential epochs and task trials. These analyses were constrained to task trials in which the letter array was presented. Additional models examined the effects of focus and task-unrelated thought ratings as predictors of microstates instead of trial accuracy, and these analyses were constrained to task trials in which the mind wandering probe questions were presented. Effects of sequential epoch and sequential trial were interpreted from models including the effects of task accuracy, but these effects are also summarized in tables from models including mind wandering as a predictor.

Effects of epoch and trial number were centered to the first 4000 msec epoch of the trial (epoch 1 = 0) and first trial of the task (trial 1 = 0), respectively. Epoch-level data (level 1) were hierarchically nested within different trial wait times (level 2) and participants (level 3). Random slopes and covariances representing trial-level and participant-level differences in change over sequential epochs were included in models. Random slopes and covariances representing participant-level differences in change over sequential task trials were also included in models. Epochs were excluded from all analyses if more than 40% of an epoch’s samples went uncategorized as microstates (< 0.03% of epochs met this criteria). Type III F-tests are reported for omnibus tests of fixed effects, and parameter estimates and simple effects are interpreted when omnibus effects are significant. To account for increased Type 1 error rates because of our use of five microstate dependent measures, we set the significance level of all Type III F-tests tests and all follow-up comparisons of model parameters in our mixed effects models to α = 0.010.

Mixed effects models were estimated with restricted maximum likelihood using the PROC HPMIXED procedure in SAS 9.4. This procedure is specifically optimized to enable mixed effects modeling in cases involving a very large number of observations or models effects. Although this procedure allows for less flexibility in modifying the estimation methods or residual error structure, its use was necessary to compute effects at the trial epoch level and test hypotheses regarding change in microstates as a function of time during the wait time delay because of the computational demand imposed by the large number of data observations and three-level data structure. Importantly, we ensured all significant effects of epoch and trial number aligned with additional sensitivity analyses, in which the overall dataset was reduced in size by averaging across epochs or trials and estimating models with restricted maximum likelihood using PROC MIXED. These analyses included both trials in which the letter array and mind wandering probes were presented to ensure effects were invariant to trial type. We interpreted the random effects from these models and evaluated the significance of random effects using loglikelihood tests of change in model fit.

#### Transition probabilities

Additionally, we examined whether the probabilities of microstates to transition from one configuration to another in the EEG time series differed as a function of the fixed effects of trial accuracy (correct vs. incorrect trials), sequential epoch number (i.e., time during the wait time delay), and sequential trial number (i.e., time-on-task) in a series of mixed effects models using PROC HPMIXED. For transition analyses, the microstate time series was transformed so that the duration of each microstate was ignored by collapsing consecutive samples of the same microstate configuration into a single observation (i.e., a Markov jump process). The set of six topographic configurations (A–F) allowed for 30 unique pairs of Markov-chain transition probabilities. Transitions of microstates to unassigned epochs were excluded from these analyses. Probabilities for each transition pair were examined in separate mixed effects models. To account for increased Type 1 error rates in these additional exploratory analyses, we set the significance level of all comparisons of model parameters in our mixed effects models to α = 0.001.

## Results

### Target Detection Accuracy and Mind Wandering Change as a Function of Trial Wait Time

The overall association between SACT task performance and mind wandering was characterized through a series of bivariate correlations. Correlations indicated that individual’s overall accuracy was moderately associated with their levels of focus (r = 0.283, p = 0.049, 95% CI [0.002, 0.523]) and task-unrelated thought (r = −0.305, p = 0.033, 95% CI [−0.540, −0.026]). Mean levels of accuracy (M = 0.659, SD = 0.136), focus (M = 3.523, SD = 0.667), and task-unrelated thought (M = 2.219, SD = 0.810), were comparable to prior work using this task (i.e., Zanesco et al., 2026b).

#### Accuracy

Growth curve models were used to characterize differences in task accuracy as a function of the linear and quadratic effects of trial wait time. Accuracy was aggregated for participants across sets of trials for each unique wait time and values were nested within individuals in analyses. We observed that accuracy decreased as a function of the linear (b = −2.3314, SE = 0.2836, p < .001, 95% CI [−2.8886, −1.7742]) and quadratic (b = 0.0317, SE = 0.0068, p < .001, 95% CI [0.0183, 0.0451]) effects of trial wait time relative to the intercept (wait time = 0 seconds) of 92.3255 (SE = 3.1239, p < .001, 95% CI [86.0445, 98.6066]). These effects indicate that participants’ average detection accuracy worsened as they had to maintain their attention over longer durations during the wait times of trials. However, the rate of decline slowed as time increased.

#### Focus

Growth curve models were used to characterize differences in focus scores as a function of the linear effects of trial wait time and sequential trial number. Focus decreased as a function of the linear effects of trial wait time (b = −0.0118, SE = 0.0041, p = .004, 95% CI [−0.0198, −0.0038]), sequential trial number (b = −0.0066, SE = 0.0029, p = .026, 95% CI [−0.0124, −0.0008]), and the interaction between wait time and trial number (b = −0.0003, SE = 0.0001, p = .008, 95% CI [−0.0005, −0.0007]), relative to the intercept of 4.1212 (SE = 0.1244, p < .001, 95% CI [3.8737, 4.3687]). These effects indicate that focus worsened to a greater degree when individuals had to maintain their attention over a long trial wait time at the end of the task relative to the beginning.

#### Task-unrelated thought

Growth curve models were used to characterize differences in task-unrelated thought (TUT) scores as a function of the linear effects of trial wait time and sequential trial number. TUT increased as a function of the linear effects of trial wait time (b = 0.0155, SE = 0.004, p < .001, 95% CI [0.0078, 0.0231]) and sequential trial number (b = 0.0093, SE = 0.0026, p < .001, 95% CI [0.0041, 0.0145]), relative to the intercept of 1.5837 (SE = 0.1102, p < .001, 95% CI [1.3646, 1.8028]). There was no interaction between wait time and trial number (b = 0.0001, SE = 0.0001, p = .469, 95% CI [−0.0001, 0.0002]). We simplified the model to estimate the effects of trial wait time and sequential trial number without the interaction term. TUT increased as a function of the linear effects of trial wait time (b = 0.0176, SE = 0.003, p < .001, 95% CI [0.0124, 0.0228]) and sequential trial number (b = 0.0105, SE = 0.0020, p < .001, 95% CI [0.0066, 0.0145]), relative to the intercept of 1.5439 (SE = 0.0956, p < .001, 95% CI [1.3518, 1.7361]). These effects indicate that TUT increased to a greater degree when individuals had to maintain their attention over a long trial wait time and as the task progressed over time.

### Microstates Differentiate Correct from Incorrect Trials

The activity and dynamics of microstates were quantified for epochs in the wait time delay of trials. We first evaluated differences in the activity and dynamics of microstates between trials in which the target was correctly detected compared to incorrectly detected. The model estimated effects of trial accuracy are summarized below for significant omnibus effects (see Table 2) for each dependent measure from mixed-effects models including the fixed effects of trial accuracy, sequential epoch, sequential trial number (i.e., time-on-task), and their two-way interactions with microstate configuration. Simple effects of trial accuracy, epoch, and trial number from these five models are summarized in Table 3 for each microstate configuration. Figure 3 depicts the model estimated means for epochs from correct and incorrect trials for all microstates, alongside the model means comparisons between correct and incorrect trials.

**Table 1.** Model estimated means for microstate.

| Measure | Mean Estimate [95% CI] |
| --- | --- |
| GEV |  |
| A | 5.7852 [5.5253, 6.0451] |
| B | 8.8619 [8.6020, 9.1218] |
| C | 29.4939 [29.2340, 29.7538] |
| D | 7.6331 [7.3732, 7.8930] |
| E | 6.1776 [5.9177, 6.4375] |
| F | 2.5301 [2.2702, 2.7900] |
| Duration |  |
| A | 63.5807 [62.7896, 64.3719] |
| B | 66.5820 [65.7914, 67.3725] |
| C | 84.7724 [83.9822, 85.5627] |
| D | 66.3532 [65.5616, 67.1447] |
| E | 65.3416 [64.5494, 66.1338] |
| F | 60.6714 [59.8775, 61.4653] |
| Occurrence |  |
| A | 1.6370 [1.5949, 1.6792] |
| B | 2.0758 [2.0337, 2.1180] |
| C | 3.4692 [3.4270, 3.5113] |
| D | 2.0747 [2.0326, 2.1169] |
| E | 1.8313 [1.7891, 1.8734] |
| F | 1.0435 [1.0014, 1.0857] |
| Num. Time Samples |  |
| A | 115.94 [114.21, 117.68] |
| B | 157.12 [155.39, 158.86] |
| C | 351.03 [349.30, 352.77] |
| D | 157.07 [155.34, 158.81] |
| E | 136.37 [134.64, 138.10] |
| F | 68.731 [66.997, 70.465] |
| GFP |  |
| A | 3.9426 [3.6092, 4.2759] |
| B | 4.1294 [3.7961, 4.4627] |
| C | 4.6454 [4.3121, 4.9787] |
| D | 3.9649 [3.6316, 4.2982] |
| E | 3.8764 [3.5431, 4.2098] |
| F | 3.6977 [3.3644, 4.0310] |

Note: Model estimates of mean microstate global explained variance (GEV), duration (in msec), occurrence rate (times per second, Hz), number time samples, and global field power (GFP in $\mu V$ ), are provided from multilevel mixed-effects models based on the fixed effects of microstate configuration from 49 participants and 14,479 epochs of EEG from the wait time delays of task trials. 95% confidence intervals (CI) accompany the model estimated means.

**Table 2.** Omnibus effects of accuracy, sequential epoch, and sequential trial on microstate dependent measures from mixed models.

| Effect | GEV |  | Duration |  | Occurrence Rate |  | Num. Time Samples |  | GFP |  |
| --- | --- | --- | --- | --- | --- | --- | --- | --- | --- | --- |
|  | *F* | *p* | *F* | *p* | *F* | *p* | *F* | *p* | *F* | *p* |
| Configuration | 3143.15 | < .001 | 403.15 | < .001 | 1482.44 | < .001 | 2065.01 | < .001 | 229.92 | < .001 |
| Accuracy | 0.22 | .636 | 0.49 | .483 | 0.50 | .480 | 0.06 | .802 | 52.98 | < .001 |
| Epoch | 0.93 | .335 | 4.23 | .040 | 4.12 | .042 | 0.30 | .581 | 2.69 | 0.101 |
| Trial | 5.19 | .023 | 1.76 | .184 | 8.50 | .004 | 0.07 | .794 | 43.19 | < .001 |
| Accuracy × Configuration | 26.65 | < .001 | 5.65 | < .001 | 4.92 | < .001 | 20.32 | < .001 | 7.17 | < .001 |
| Epoch × Configuration | 2.16 | .055 | 1.54 | .173 | 3.96 | .001 | 3.97 | .001 | 1.45 | 0.202 |
| Trial × Configuration | 23.68 | < .001 | 2.06 | .067 | 3.14 | .008 | 12.17 | < .001 | 9.63 | < .001 |

Note: The omnibus results of Type III *F* tests are provided from mixed models of the fixed effects of trial accuracy, sequential epoch, and sequential trial number, and their two-way interactions for dependent measures of mean microstate global explained variance (GEV), duration (in msec), occurrence rate (times per second, Hz), number time samples, and global field power (GFP in μV).

**Table 3.** Simple effects of accuracy, sequential epoch, and sequential trial on microstate dependent measures from mixed models.

| Effect | GEV (%) | Duration (msec) | Occurrence Rate (Hz) | Num. Time Samples | GFP (μV) |
| --- | --- | --- | --- | --- | --- |
| Accuracy (Map A) | 0.1125 (0.1333) | 0.3333 (0.3448) | -0.0035 (0.0161) | 0.7950 (1.6592) | -0.0501 (0.0164)** |
| Accuracy (Map B) | 0.1552 (0.1333) | 0.0702 (0.3430) | 0.0159 (0.0161) | 2.3379 (1.6592) | -0.0822 (0.0164)*** |
| Accuracy (Map C) | -1.4038 (0.1333)*** | -1.7053 (0.3419)*** | -0.0532 (0.0161)*** | -14.9864 (1.6592)*** | -0.1225 (0.0163)*** |
| Accuracy (Map D) | 0.2879 (0.1333)* | 0.4275 (0.3455) | -0.0103 (0.0161) | 1.6961 (1.6592) | -0.0589 (0.0165)*** |
| Accuracy (Map E) | 0.3912 (0.1333)** | 0.3009 (0.3474) | 0.0422 (0.0161)** | 5.3613 (1.6592)** | -0.0045 (0.0166) |
| Accuracy (Map F) | 0.2957 (0.1333)* | -0.0576 (0.3535) | 0.0381 (0.0161)* | 3.7467 (1.6592)* | -0.0203 (0.0168) |
| Epoch (Map A) | 0.0475 (0.0270) | 0.0043 (0.0736) | 0.0062 (0.0033) | 0.5618 (0.3354) | 0.0132 (0.0066)* |
| Epoch (Map B) | -0.0146 (0.0270) | -0.0494 (0.0732) | -0.0006 (0.0033) | -0.1361 (0.3354) | 0.0065 (0.0066) |
| Epoch (Map C) | -0.0306 (0.0270) | -0.2384 (0.0730)** | -0.0001 (0.0033) | -0.8084 (0.3354)* | 0.0080 (0.0066) |
| Epoch (Map D) | -0.0130 (0.0270) | -0.0864 (0.0737) | -0.0027 (0.0033) | -0.3211 (0.3354) | 0.0049 (0.0066) |
| Epoch (Map E) | 0.0699 (0.0270)** | -0.0751 (0.0741) | 0.0147 (0.0033)*** | 1.0801 (0.3354)** | 0.0149 (0.0066)* |
| Epoch (Map F) | 0.0064 (0.0270) | -0.0215 (0.0753) | 0.0004 (0.0033) | 0.0804 (0.3354) | 0.0108 (0.0066) |
| Trial (Map A) | -0.0038 (0.0040) | 0.0025 (0.0112) | -0.0013 (0.0005)* | -0.0907 (0.0493) | 0.0091 (0.0014)*** |
| Trial (Map B) | -0.0006 (0.0040) | 0.0068 (0.0111) | -0.0008 (0.0005) | -0.0022 (0.0493) | 0.0091 (0.0014)*** |
| Trial (Map C) | 0.0420 (0.0040)*** | 0.0361 (0.0111)** | 0.0008 (0.0005) | 0.3389 (0.0493)*** | 0.0083 (0.0014)*** |
| Trial (Map D) | -0.0055 (0.0040) | -0.0036 (0.0112) | -0.0010 (0.0005)* | -0.0675 (0.0493) | 0.0082 (0.0014)*** |
| Trial (Map E) | -0.0019 (0.0040) | 0.0024 (0.0113) | -0.0012 (0.0005)* | -0.0670 (0.0493) | 0.0075 (0.0014)*** |
| Trial (Map F) | -0.0044 (0.0040) | 0.0095 (0.0114) | -0.0014 (0.0005)** | -0.0750 (0.0493) | 0.0116 (0.0014)*** |

Note: Model parameters representing simple effects of accuracy (correct vs. incorrect), sequential epoch, and sequential trial on microstate dependent measures for configurations A through F from mixed-effects models. Simple effects are organized in columns for each dependent measure. \* $p < .05$ , \*\* $p < .01$ , \*\*\* $p < .001$

**Figure 3.**
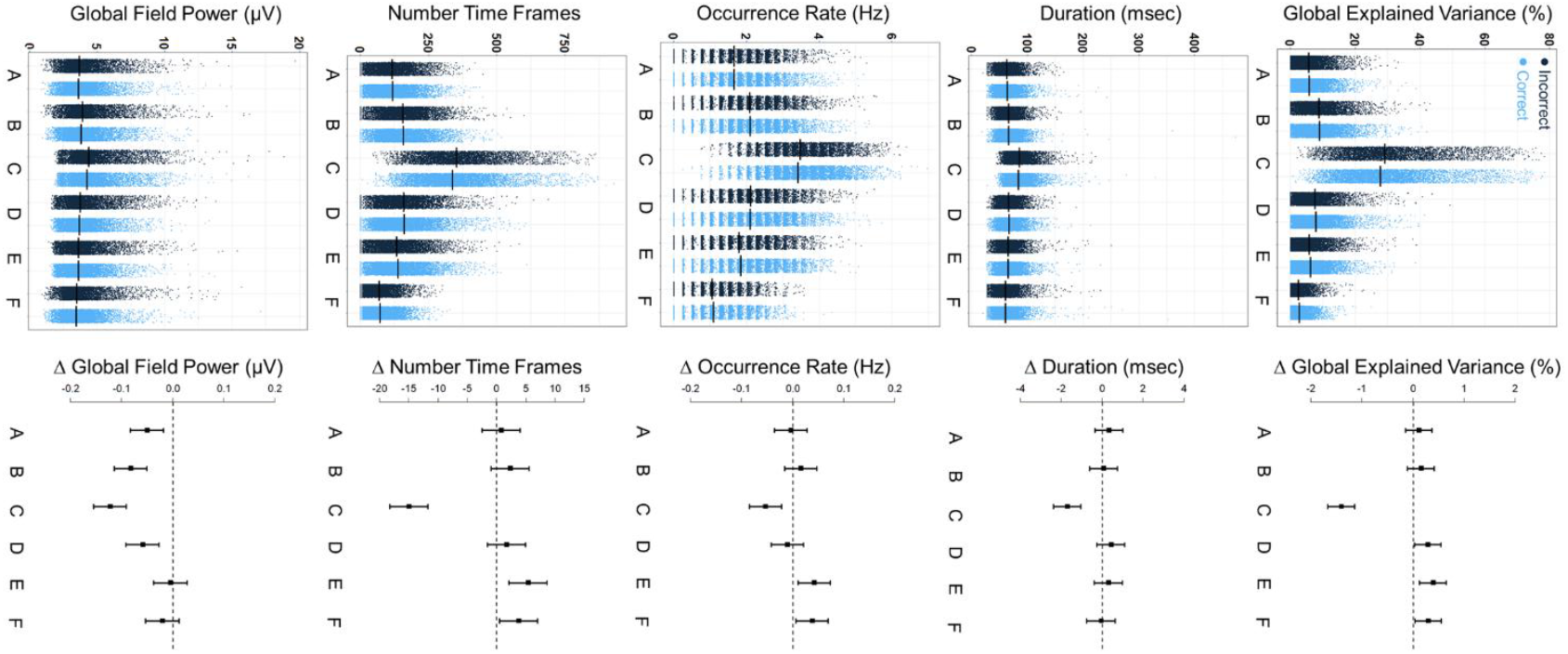
Observed data are shown on the left for epochs from trials in which individuals correctly detected the target stimuli in light blue and from incorrect trials in dark blue for dependent measures of microstates. Model estimated means are provided as black bars. The model estimated difference (Δ) between correct and incorrect trials and 95% confidence intervals are shown on the right for each microstate.

We found that the dynamics of microstates C and E differentiated correctly detected trials from incorrect trials. Microstate configuration C explained significantly less GEV (b = −1.4038, SE = 0.1333, p < .001, 95% CI [−1.6650, −1.1426]), had shorter duration (b = −1.7053 msec, SE = 0.3419, p < .001, 95% CI [−2.3754, −1.0352]), occurred less frequently (b = −0.0532 Hz, SE = 0.0161, p = .001, 95% CI [−0.0848, −0.0216]), and occupied fewer total time samples (b = −14.9864, SE = 1.6592, p < .001, 95% CI [−18.2384, −11.7344]), on epochs when individuals correctly detected the target stimuli amid the letter array relative to incorrect trials. In contrast, configuration E explained significantly more GEV (b = 0.3912, SE = 0.1333, p = .003, 95% CI [0.1300, 0.6524]), occurred more frequently (b = 0.0422 Hz, SE = 0.0161, p = .009, 95% CI [0.0105, 0.0738]), and occupied more total time samples (b = 5.3613, SE = 1.6592, p = .001, 95% CI [2.1092, 8.6133]) for epochs of correct trials relative to incorrect trials.

In addition, the overall amplitude of several microstates differentiated correctly detected trials from incorrect trials. Microstate configuration A (b = −0.0501 uV, SE = 0.0164, p = .002, 95% CI [−0.0823, −0.0179]), configuration B (b = −0.0822 uV, SE = 0.0164, p < .001, 95% CI [−0.1142, −0.0501]), configuration C (b = −0.1225 uV, SE = 0.0164, p < .001, 95% CI [−0.1545, −0.0905]), and configuration D (b = −0.0589 uV, SE = 0.0164, p < .001, 95% CI [−0.0912, −0.0266]), all had significantly lower GFP on epochs when individuals correctly detected the target stimuli amid the letter array relative to incorrect trials.

### Microstate Dynamics Change as Attention is Maintained Over the Wait Time Delay

We next evaluated change in microstates as a function of sequential epochs of the wait time delay. The model estimated effects of epochs are summarized below for significant omnibus effects (see Table 2) for each dependent measure from mixed-effects models including the fixed effects of trial accuracy, sequential epoch, sequential trial number (i.e., time-on-task), and their two-way interactions with microstate configuration. Simple effects from models are summarized in Table 3. The dynamics of microstate E were observed to change over time as a function of sequential epochs in the wait time delay. Microstate configuration E occurred significantly more frequently (b = 0.0147 Hz, SE = 0.0033, p < .001, 95% CI [0.0082, 0.0212]) and occupied more time samples (b = 1.0801, SE = 0.3354, p = .001, 95% CI [0.4226, 1.7375]), each sequential epoch as individuals attempted to maintain their attention over the wait time of trials. Microstate E was estimated to occur 0.1324 (SE = 0.0298, p < .001, 95% CI [0.0740, 0.1907]) more times per second and occupy 9.7207 (SE = 3.0189, p = .001, 95% CI [3.8035, 15.6378]) more total time samples in the epochs, at the end of 40 second trials compared to the first epoch at the beginning of trials.

We also conducted sensitivity analyses to confirm these longitudinal trends for microstate E and evaluate the extent of interindividual differences in intraindividual change during the wait time delay of trials. Measures of microstate E were aggregated across trials for each sequential epoch and data were nested within individuals for analysis of the effects of epoch using linear mixed-effects models. Microstate E occurred more frequently (b = 0.0098 Hz, SE = 0.0031, p = .003, 95% CI [0.0035, 0.0161]) and occupied more total time samples (b = 0.7268, SE = 0.2856, p = .014, 95% CI [0.1525, 1.3011]) each sequential epoch, as individuals attempted to maintain their attention over the wait time delay of trials. Figure 4 depicts the observed means for microstate E aggregated across trials for epochs alongside model estimates of effects. The random effects for the slope and covariance of epochs were also significant for occurrence rate, −2ΔLL(2) = 15, p < .001, and number of time samples, −2ΔLL(2) = 18.3, p < .001, suggesting interindividual variability in rates of change over the wait time delay of trials. Random effects CIs suggested that 95% of participants demonstrated a per-epoch linear change in occurrence rate of between −0.0216 and 0.0412 and a per-epoch linear change in the number of time samples between −2.1689 and 3.6225.

**Figure 4.**
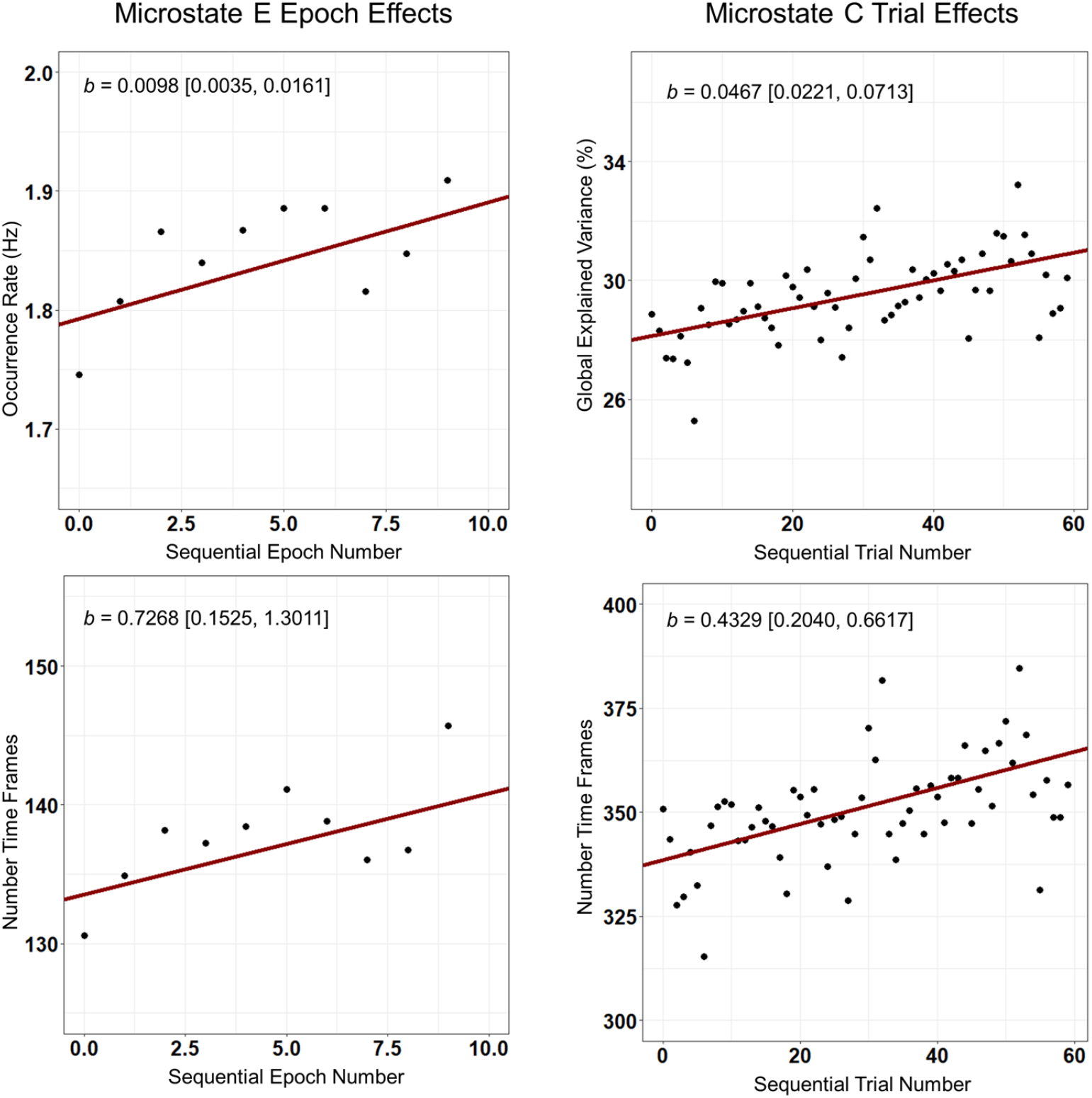
Growth curve model estimated trajectories for microstate E (left) and microstate C (right) are plotted in red as a function of sequential 4000 msec epochs and sequential task trials. Effects of epoch and trial number were centered to the first 4000 msec epoch of the trial (epoch 1 = 0) and first trial of the task (trial 1 = 0), respectively. The observed means for each sequential epoch and sequential trial shown as black dots. Slope estimates are also provided alongside 95% confidence intervals.

### Microstate Dynamics Change Over the Task Session

We next evaluated changes in microstates as a function of sequential trials over the task session. The model estimated effects of sequential trials are summarized below for significant omnibus effects (see Table 2) for each dependent measure from mixed-effects models including the fixed effects of trial accuracy, sequential epoch, sequential trial number (i.e., time-on-task), and their two-way interactions with microstate configuration. Simple effects are summarized in Table 3. Microstate configuration C was more prevalent as the task progressed over trials. Microstate configuration C explained significantly more GEV (b = 0.0420, SE = 0.0040, p < .001, 95% CI [0.0342, 0.0497]) and occupied more total time samples (b = 0.3389, SE = 0.0493, p < .001, 95% CI [0.2423, 0.4354]) as the task progressed in time over sequential task trials. Microstate C was therefore estimated to explain 2.4756 (SE = 0.2334, p < .001, 95% CI [2.0181, 2.9331]) more units of GEV and occupy 19.99 (SE = 2.9064, p < .001, 95% CI [14.2978, 25.6910]) more total time samples at the end of the task on the 60th trial compared to the beginning. In addition, microstate configuration F occurred less frequently (b = −0.0014 Hz, SE = 0.0005, p = .006, 95% CI [−0.0024, −0.0004]) over sequential task trials.

Finally, the amplitude of all microstates on average increased as the task progressed over trials: GFP increased for microstate configuration A (b = 0.0091 uV, SE = 0.0014, p < .001, 95% CI [0.0063, 0.0119]), configuration B (b = 0.0091 uV, SE = 0.0014, p < .001, 95% CI [0.0063, 0.0119]), configuration C (b = 0.0116 uV, SE = 0.0014, p < .001, 95% CI [0.0088, 0.0144]), configuration D (b = 0.0083 uV, SE = 0.0014, p < .001, 95% CI [0.0055, 0.0112]), configuration E (b = 0.0082 uV, SE = 0.0014, p < .001, 95% CI [0.0054, 0.0110]), and configuration F (b = 0.0075 uV, SE = 0.0014, p < .001, 95% CI [0.0047, 0.0103]). While all microstates increased in amplitude over sequential trials, microstate C increased to the greatest extent.

We also conducted sensitivity analyses to confirm these longitudinal trends for microstate C and evaluate the extent of interindividual differences in intraindividual change over the task session. Measures of microstate C were aggregated for participants across epochs for trials and data were nested within individuals for analysis of the effects of sequential trial using linear mixed-effects models. Microstate configuration C was more prevalent as the task progressed over trials. Microstate configuration C explained significantly more GEV (b = 0.0467, SE = 0.0122, p < .001, 95% CI [0.0221, 0.0713]) and occupied more total time samples (b = 0.4329, SE = 0.1138, p < .001, 95% CI [0.2040, 0.6617]) as the task progressed in time over sequential task trials. Figure 4 depicts the observed means for microstate C aggregated across epochs for trials alongside model estimates of effects. The random effects for the slope and covariance of sequential trials were also significant for GEV, −2ΔLL(2) = 97.4, p < .001, and number of time samples, −2ΔLL(2) = 76.8, p < .001, suggesting interindividual variability in change over the task session. Random effects CIs suggested that 95% of participants demonstrated a per-trial linear change in GEV of microstate C between −0.1023 and 0.1957 and a per-trial linear change in the number of time samples between −0.9240 and 1.7898.

### Wait Times of Previous Trials Influence Microstates on Subsequent Trials

We evaluated the effects of previous trial (trial T – 1) wait time on current trial microstates by including the effects of previous trial wait time in additional mixed-effects models, alongside the effects of trial accuracy, epoch, and sequential trial number. The first task trial was included in analyses with a previous trial wait time equal to 0. We observed a significant interaction between previous trial wait time and microstate configuration for models of microstate GEV, F(5, 64314) = 10.49, p < .001, occurrence rate, F(5, 64314) = 3.27, p = .006, and number of time samples, F(5, 64314) = 7.09, p < .001. There was also a significant interaction for microstate duration, F(5, 62805) = 3.29, p = .006, but none of the simple effects were significant (ps > .01).

Microstate configuration C was more prevalent on the current trial when the previous trial had a longer wait time delay. Microstate configuration C explained significantly more GEV (b = 0.0352, SE = 0.0053, p < .001, 95% CI [0.0248, 0.0456]), occurred more frequently (b = 0.0021 Hz, SE = 0.0006, p = .001, 95% CI [0.0008, 0.0033]), and occupied more total time samples (b = 0.3401, SE = 0.0663, p < .001, 95% CI [0.2101, 0.4701]) for each additional second of the length of the wait time in the previous trial. Microstate C was therefore estimated to explain 1.4083 (SE = 0.2126, p < .001, 95% CI [0.9916, 1.8251]) more units of GEV, occur 0.0833 (SE = 0.0257, p = .001, 95% CI [0.0329, 0.1338]) more times per second, and occupy 13.6036 (SE = 2.6525, p < .001, 95% CI [8.4048, 18.8024]) more time samples on the current trial when the previous trial was 40 seconds in duration. Finally, there was also a significant main effect of previous trial wait time for microstate GFP, F(1, 62805) = 51.70, p < .001, but no significant interaction with configuration, F(5, 62805) = 2.02, p = .072. We simplified the model by removing the interaction term to estimate this main effect. GFP was greater for all configurations on average (b = 0.0022 µV, SE = 0.0003, p < .001, 95% CI [0.0016, 0.0028]) for each additional second of duration of the wait time in the previous trial.

### Microstates Are Associated with Focus and Task-Unrelated Thought Ratings

We examined the effects of mind wandering (focus and task-unrelated thought), sequential epoch, sequential trial number (i.e., time-on-task), and their two-way interactions with microstate configuration, separately for focus, task-unrelated thought, and each microstate dependent measure. The model estimated effects are described for significant omnibus effects (see Table 4). Simple effects from models including focus ratings are summarized in Table 5 for each microstate configuration, whereas effects from models including task-unrelated thought ratings are summarized in Table 6. Figure 5 depicts the model estimated effects for all microstates.

**Table 4.** Omnibus effects of accuracy, sequential epoch, and sequential trial on microstate dependent measures from mixed models.

| Effect | GEV |  | Duration |  | Occurrence Rate |  | Num. Time Samples |  | GFP |  |
| --- | --- | --- | --- | --- | --- | --- | --- | --- | --- | --- |
|  | *F* | *p* | *F* | *p* | *F* | *p* | *F* | *p* | *F* | *p* |
| Focus |  |  |  |  |  |  |  |  |  |  |
| Configuration | 371.94 | < .001 | 39.48 | < .001 | 200.68 | < .001 | 241.62 | < .001 | 29.47 | < .001 |
| Focus | 0.02 | .895 | 0.51 | .477 | 0.19 | .663 | 0.12 | .724 | 0.13 | 0.723 |
| Epoch | 0.24 | .626 | 7.74 | .005 | 1.72 | .189 | 0.08 | .782 | 6.70 | 0.01 |
| Trial | 2.25 | .134 | 3.79 | .052 | 0.90 | .342 | 0.01 | .972 | 26.65 | < .001 |
| Focus × Configuration | 4.22 | < .001 | 1.09 | .366 | 10.72 | < .001 | 5.07 | < .001 | 1.17 | 0.321 |
| Epoch × Configuration | 1.49 | .191 | 0.10 | .992 | 1.88 | .095 | 1.85 | .100 | 0.96 | 0.441 |
| Trial × Configuration | 4.90 | < .001 | 1.36 | .238 | 1.77 | .116 | 3.83 | .002 | 1.96 | 0.081 |
| Task-unrelated thought |  |  |  |  |  |  |  |  |  |  |
| Configuration | 780.44 | < .001 | 97.45 | < .001 | 353.60 | < .001 | 497.49 | < .001 | 50.49 | < .001 |
| TUT | 0.06 | .814 | 1.81 | .179 | 0.01 | .933 | 0.01 | .950 | 16.53 | < .001 |
| Epoch | 0.22 | .638 | 8.41 | .004 | 1.98 | .159 | 0.06 | .814 | 5.95 | 0.015 |
| Trial | 2.26 | .133 | 4.49 | .034 | 0.65 | .421 | 0.01 | .963 | 23.43 | < .001 |
| TUT × Configuration | 7.87 | < .001 | 0.22 | .952 | 28.33 | < .001 | 10.96 | < .001 | 1.89 | 0.092 |
| Epoch × Configuration | 1.37 | .232 | 0.15 | .981 | 1.99 | .077 | 1.68 | .136 | 0.91 | 0.471 |
| Trial × Configuration | 5.92 | < .001 | 1.96 | .082 | 2.28 | .044 | 5.05 | < .001 | 2.03 | 0.072 |

Note: The omnibus results of Type III *F* tests are provided from mixed models of the fixed effects of focus or task-unrelated thought, sequential epoch, and sequential trial number, and their two-way interactions for dependent measures of mean microstate global explained variance (GEV), duration (in msec), occurrence rate (times per second, Hz), number time samples, and global field power (GFP in μV).

**Table 5.**
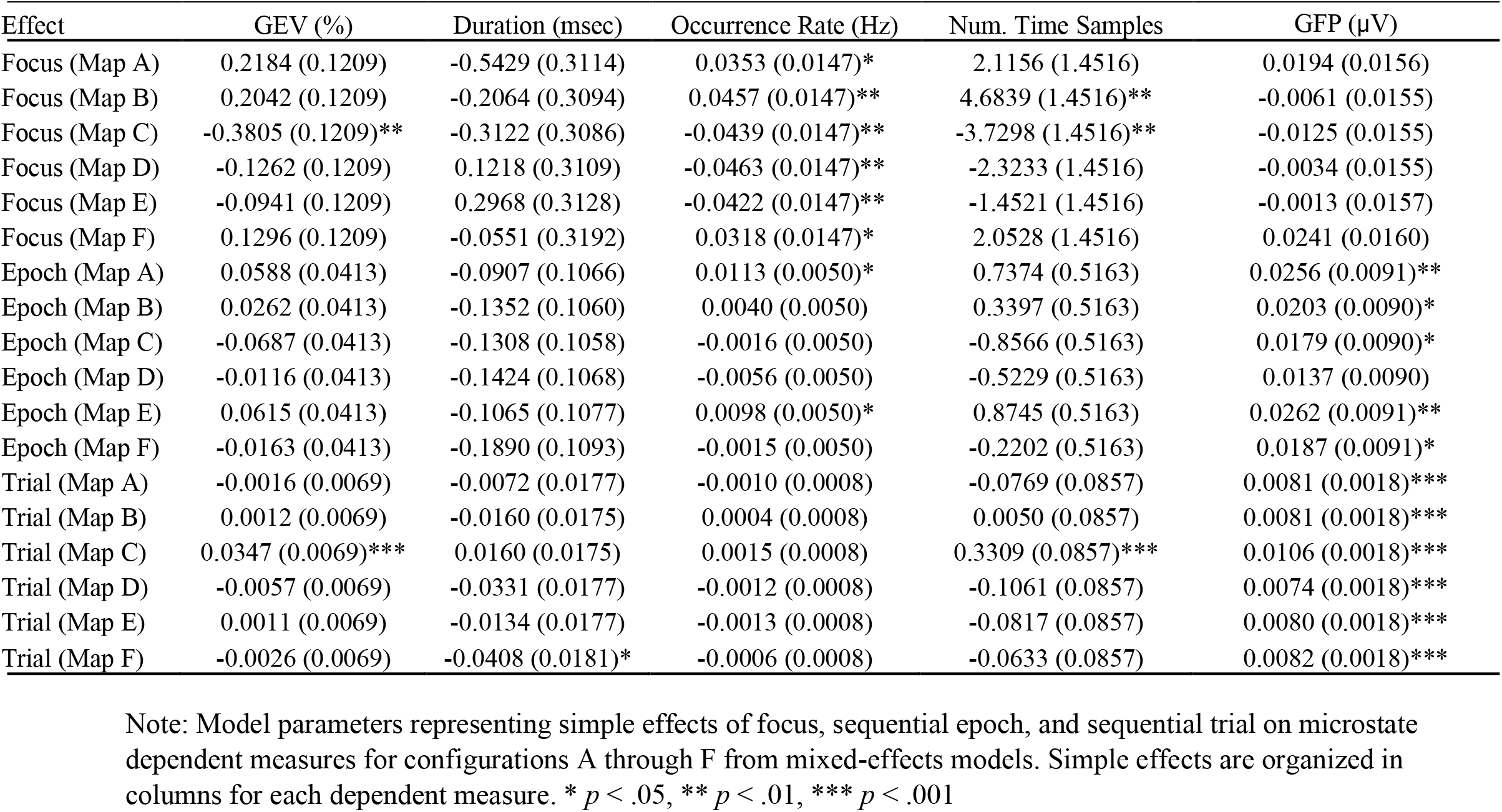
Simple effects of focus, sequential epoch, and sequential trial on microstate dependent measures from mixed models.

| Effect | GEV (%) | Duration (msec) | Occurrence Rate (Hz) | Num. Time Samples | GFP (μV) |
| --- | --- | --- | --- | --- | --- |
| Focus (Map A) | 0.2184 (0.1209) | -0.5429 (0.3114) | 0.0353 (0.0147)* | 2.1156 (1.4516) | 0.0194 (0.0156) |
| Focus (Map B) | 0.2042 (0.1209) | -0.2064 (0.3094) | 0.0457 (0.0147)** | 4.6839 (1.4516)** | -0.0061 (0.0155) |
| Focus (Map C) | -0.3805 (0.1209)** | -0.3122 (0.3086) | -0.0439 (0.0147)** | -3.7298 (1.4516)** | -0.0125 (0.0155) |
| Focus (Map D) | -0.1262 (0.1209) | 0.1218 (0.3109) | -0.0463 (0.0147)** | -2.3233 (1.4516) | -0.0034 (0.0155) |
| Focus (Map E) | -0.0941 (0.1209) | 0.2968 (0.3128) | -0.0422 (0.0147)** | -1.4521 (1.4516) | -0.0013 (0.0157) |
| Focus (Map F) | 0.1296 (0.1209) | -0.0551 (0.3192) | 0.0318 (0.0147)* | 2.0528 (1.4516) | 0.0241 (0.0160) |
| Epoch (Map A) | 0.0588 (0.0413) | -0.0907 (0.1066) | 0.0113 (0.0050)* | 0.7374 (0.5163) | 0.0256 (0.0091)** |
| Epoch (Map B) | 0.0262 (0.0413) | -0.1352 (0.1060) | 0.0040 (0.0050) | 0.3397 (0.5163) | 0.0203 (0.0090)* |
| Epoch (Map C) | -0.0687 (0.0413) | -0.1308 (0.1058) | -0.0016 (0.0050) | -0.8566 (0.5163) | 0.0179 (0.0090)* |
| Epoch (Map D) | -0.0116 (0.0413) | -0.1424 (0.1068) | -0.0056 (0.0050) | -0.5229 (0.5163) | 0.0137 (0.0090) |
| Epoch (Map E) | 0.0615 (0.0413) | -0.1065 (0.1077) | 0.0098 (0.0050)* | 0.8745 (0.5163) | 0.0262 (0.0091)** |
| Epoch (Map F) | -0.0163 (0.0413) | -0.1890 (0.1093) | -0.0015 (0.0050) | -0.2202 (0.5163) | 0.0187 (0.0091)* |
| Trial (Map A) | -0.0016 (0.0069) | -0.0072 (0.0177) | -0.0010 (0.0008) | -0.0769 (0.0857) | 0.0081 (0.0018)*** |
| Trial (Map B) | 0.0012 (0.0069) | -0.0160 (0.0175) | 0.0004 (0.0008) | 0.0050 (0.0857) | 0.0081 (0.0018)*** |
| Trial (Map C) | 0.0347 (0.0069)*** | 0.0160 (0.0175) | 0.0015 (0.0008) | 0.3309 (0.0857)*** | 0.0106 (0.0018)*** |
| Trial (Map D) | -0.0057 (0.0069) | -0.0331 (0.0177) | -0.0012 (0.0008) | -0.1061 (0.0857) | 0.0074 (0.0018)*** |
| Trial (Map E) | 0.0011 (0.0069) | -0.0134 (0.0177) | -0.0013 (0.0008) | -0.0817 (0.0857) | 0.0080 (0.0018)*** |
| Trial (Map F) | -0.0026 (0.0069) | -0.0408 (0.0181)* | -0.0006 (0.0008) | -0.0633 (0.0857) | 0.0082 (0.0018)*** |

Note: Model parameters representing simple effects of focus, sequential epoch, and sequential trial on microstate dependent measures for configurations A through F from mixed-effects models. Simple effects are organized in columns for each dependent measure. \* $p < .05$ , \*\* $p < .01$ , \*\*\* $p < .001$

**Table 6.**
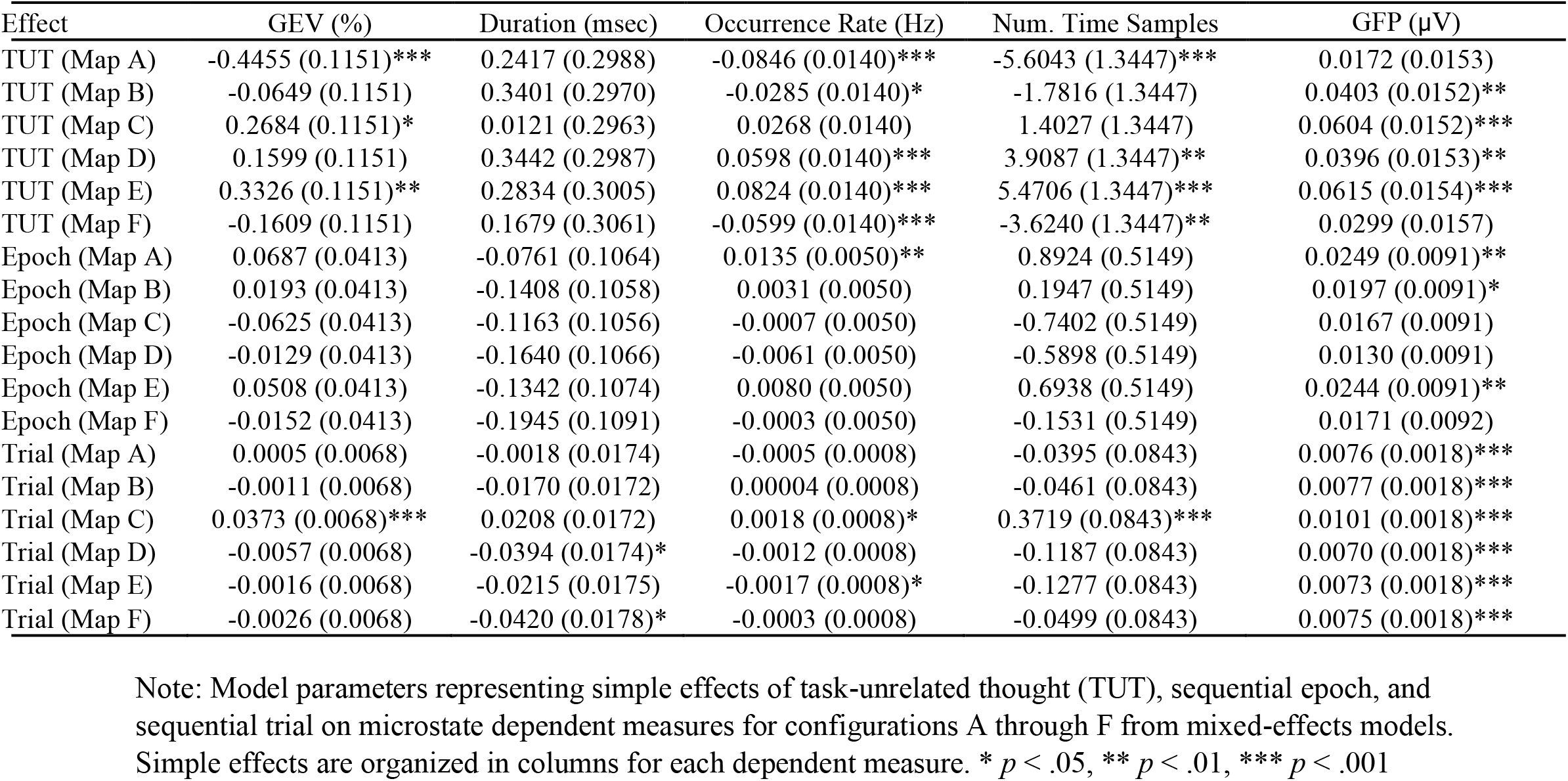
Simple effects of TUT, sequential epoch, and sequential trial on microstate dependent measures from mixed models.

**Figure 5.**
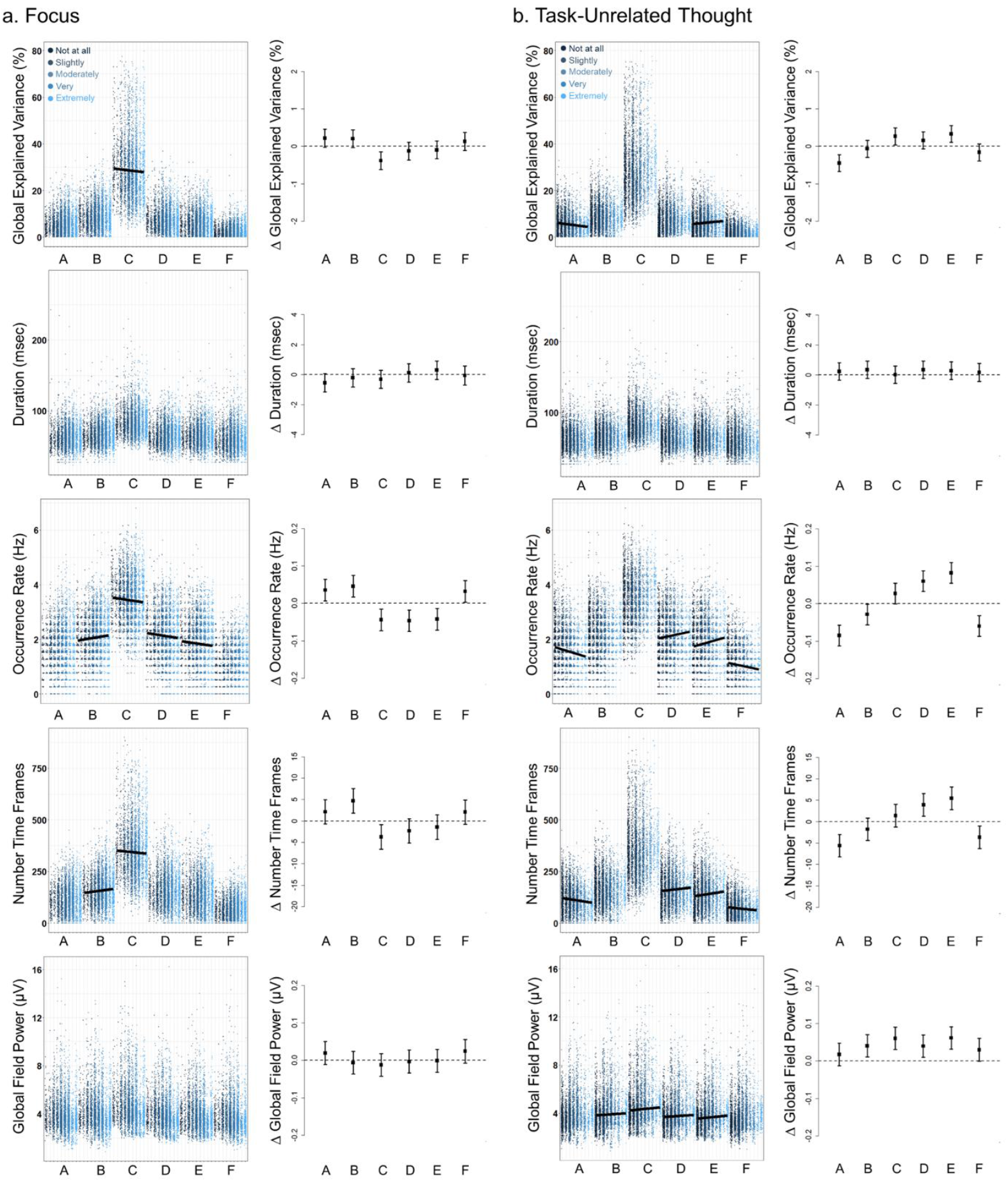
Observed data are shown for dependent measures of microstates plotted according to (a) focus or (b) task-unrelated thought ratings provided by individuals for epochs from trials with mind wandering probes. The model estimated intercept and slope describing the association between mind wandering ratings and microstates are plotted on top of the observed data for significant effects. The slope estimates and 95% confidence intervals are shown on the right of the observed data for each microstate.

#### Focus

Microstate configuration C explained significantly less GEV (b = −0.3805, SE = 0.1209, p = .002, 95% CI [−0.6175, −0.1436]), occurred less frequently (b = −0.0439, SE = 0.0147, p = .003, 95% CI [−0.0726, −0.0152]), and occupied fewer time samples (b = −3.7298, SE = 1.4516, p = .010, 95% CI [−6.5750, −0.8845]) when individuals reported being more focused on trials. On the other hand, microstate configuration B occurred significantly more frequently (b = 0.0457, SE = 0.0147, p = .002, 95% CI [0.0169, 0.0744]) and occupied more time samples (b = 4.6839, SE = 1.4516, p = .001, 95% CI [1.8386, 7.5291]) on epochs when individuals reported being more focused on trials. In addition, microstate configuration D (b = −0.0463, SE = 0.0147, p = .002, 95% CI [−0.0751, −0.0176]) and configuration E (b = −0.0422, SE = 0.0147, p = .004, 95% CI [−0.0709, −0.0135]) occurred less frequently when individuals reported being more focused.

#### Task-unrelated thought

Microstate configuration A explained less GEV (b = −0.4455, SE = 0.1151, p < .001, 95% CI [−0.6712, −0.2198]), occurred less frequently (b = −0.0846, SE = 0.0140, p < .001, 95% CI [−0.1121, −0.0570]), and occupied less time samples (b = −5.6043, SE = 1.3447, p < .001, 95% CI [−8.2401, −2.9685]) on epochs when individuals reported experiencing more task-unrelated thought. In contrast, microstate configuration D occurred more frequently (b = 0.0598, SE = 0.0140, p < .001, 95% CI [0.0323, 0.0873]) and occupied more time samples (b = 3.9087, SE = 1.3447, p = .004, 95% CI [1.2729, 6.5445]) when individuals reported experiencing more task-unrelated thought. Similarly, microstate configuration E explained more GEV (b = 0.3326, SE = 0.1151, p = .004, 95% CI [0.1069, 0.5583]), occurred more frequently (b = 0.0824, SE = 0.0140, p < .001, 95% CI [0.0549, 0.1099]), and occupied more time samples (b = 5.4706, SE = 1.3447, p < .001, 95% CI [2.8349, 8.1064]) when individuals reported experiencing more task-unrelated thought. Microstate configuration F occurred less frequently (b = −0.0599, SE = 0.0140, p < .001, 95% CI [−0.0874, −0.0324]) and occupied less time samples (b = −3.6240, SE = 1.3447, p = .007, 95% CI [−6.2597, −0.9882]) when individuals reported experiencing more task-unrelated thought. Finally, we observed a significant main effect of task-unrelated thought on microstate GFP and simplified the model by removing the interaction term to estimate this main effect. GFP increased for all configurations on average the more individuals reported experiencing task-unrelated thought (b = 0.0418 µV, SE = 0.0102, p < .001, 95% CI [0.0218, 0.0618]).

### Microstate Transition Probabilities

In a final analysis of microstate dynamics, we examined whether the probability of transitioning from one microstate configuration to another differed as a function of trial accuracy and mind wandering and changed over the wait time delay and sequential trials of the task. We first compared differences in transition probabilities of correct and incorrect trials in a series of mixed models. Separately, we examined associations between mind wandering ratings and transition probabilities. Figure 6 summarizes the results of analyses. Estimates are provided for effects representing the difference between correct and incorrect trials, change over epochs of the wait time delay, and change over sequential trials of the task. These effects result from analyses constrained to task trials in which the letter array was presented. Additionally, the effects of focus and task-unrelated thought on transition probabilities are provided. One set of transition probabilities changed significantly (p < .001) as a function of epochs of the wait time delay. Transitions from microstate B → E (b = 0.0024, SE = 0.0006, p < .001, 95% CI [0.0012, 0.0035]) increased in frequency over time during the wait time delay.

**Figure 6.**
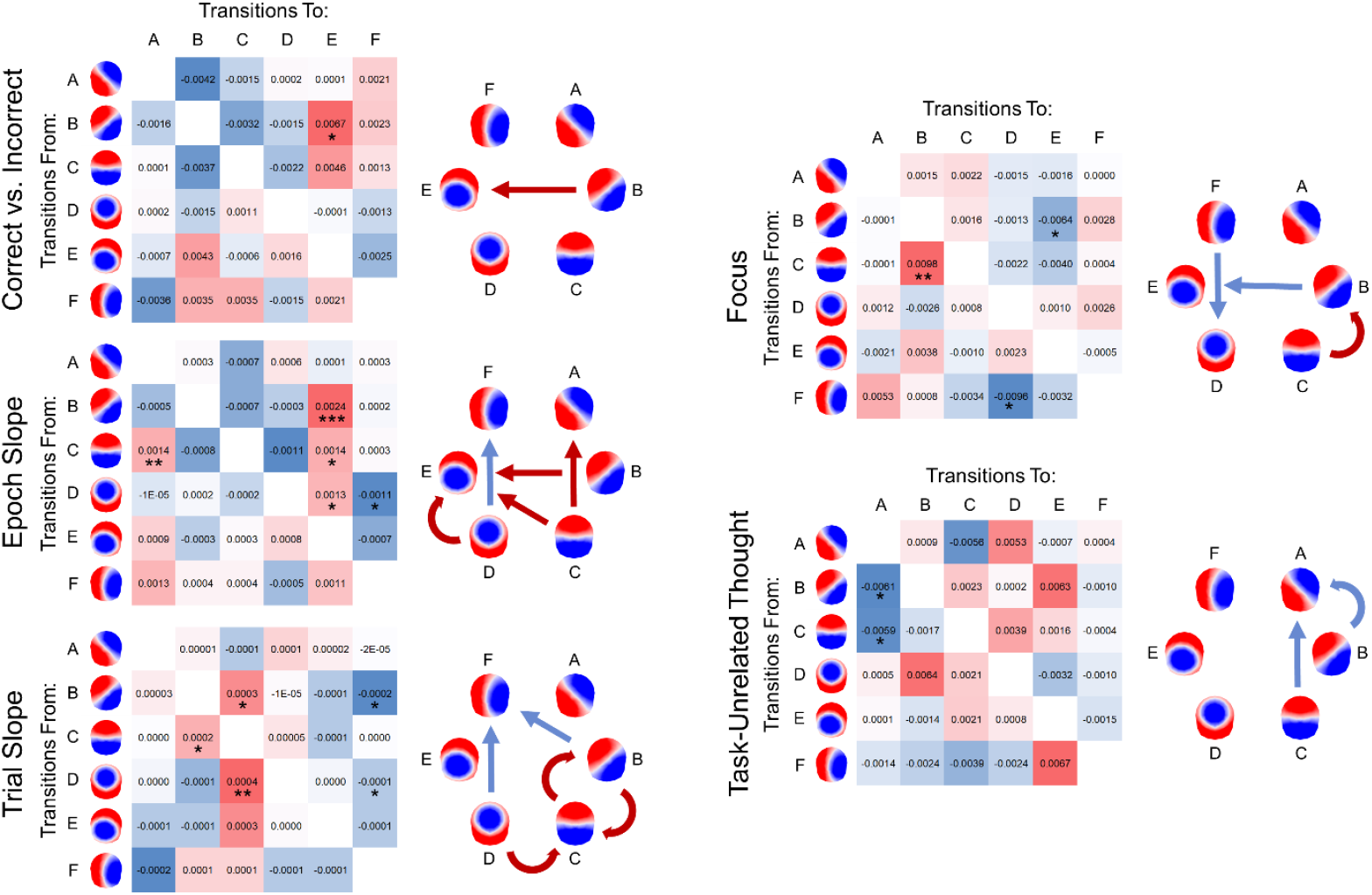
Mixed-effects model estimates of mean microstate transition probabilities are provided for all transition pairs. Transitions to unlabeled epochs of EEG are omitted. Positive associations between transition probabilities for pairs of microstate configurations are indicated by red arrows, while negative associations are indicated by blue arrows. * *p* < .05, ** *p* < .01, *** *p* < .001

## Discussion

We investigated whether the strength and temporal dynamics of distinct electroencephalographic brain states differentiated focus from inattention and changed over time as a function of the maintenance of endogenous attention during the wait time delay of task trials or over the course of the entire task session. Task trials in which individuals maintained their focus and correctly detected the target stimulus were distinguished from incorrect trials by the overall prevalence and activation strength of specific EEG microstates, particularly microstates C and E, occurring in the wait time delay of trials. The same microstates implicated in states of task focus and inattention also changed over time during the wait time delay of trials or over the course of the task session. Taken together, these findings help clarify the functional relevance of EEG microstates for human cognition and support the notion that distinct brain states and their millisecond dynamics contribute to the maintenance of attention over time and mark the occurrence of lapses of attention and episodes of mind wandering.

Notably, our findings replicate patterns of associations connecting the activity and dynamics of microstate C to states of inattention and microstate E to states of focus. We found that microstate C explained more topographic variance (GEV), occupied more of epochs’ time series, and occurred more frequently when individuals were unfocused on trials and incorrectly detected the presentation of the target stimulus. In contrast, microstate E explained more GEV, occupied more of epochs’ time series, and occurred more frequently on correct trials, relative to incorrect trials. These patterns replicate prior research linking greater prevalence of microstate C in the pre-stimulus periods of trials to probe-caught reports of mind wandering and microstate E to reports of on-task focus (Zanesco et al., 2021). Moreover, microstate C has been previously associated with behavioral indicators of inattention, including slower (Di Muccio et al., 2023) and more variable response times (Zanesco et al., 2021), whereas microstate E is associated with faster response times. Finally, a recent study found that microstate C was less prevalent and microstate E more prevalent during breath-focused attention compared to task-unconstrained periods of rest (Ngo et al., 2026). While some prior evidence has linked microstate D with attention and executive functions (Tarailis et al., 2024), we found no systematic association between the dynamics of this microstate and states of focus in our study.

The association of microstates C and E with behavioral and self-reported indicators of inattention and focus, respectively, parallels antagonistic interactions between task-positive brain networks active in support of externally directed attention, comprising the dorsal attention and frontoparietal control networks, and suppression of the default mode network (Raichle, 2015; Menon, 2023). Previous studies demonstrate that slower and more variable response times are associated with weaker anti-correlations between activity of task-positive networks and the default mode network (Kelly et al., 2008; Seeburger et al., 2024), as well as weaker task-related suppression of activity in the default mode network (Weissman et al., 2006). The occurrence of mind wandering during cognitive-behavioral tasks is also associated with increased activity in the default mode network (Christoff et al., 2009; Fox et al., 2015; Kucyi et al., 2024; Mason et al., 2007; Zhou et al., 2018). Our findings align with these neuroimaging perspectives and support the notion that the activity and dynamics of EEG microstates may result from millisecond electrophysiological interactions between and within these whole-brain neuronal networks.

Source localization of the distributed brain regions contributing to microstate C has previously identified generators in core regions of the default mode network (Bréchet et al., 2019; Custo et al., 2017) and in the medial temporal lobe subsystem of the default mode network (Ngo et al., 2026; Valt et al., 2024; Tarailis et al., 2025; Zanesco et al., 2026a). The brain generators of microstate E, however, have been less consistently investigated; sources have been identified in the anterior cingulate and salience network (Custo et al., 2017), as well as in frontal and parietal regions with functions ascribed to attention, perception, and cognitive control (Bréchet et al., 2019; Ngo et al., 2026; Zanesco et al., 2026a). Source localization of the microstate-categorized EEG in our study was tentatively consistent with the perspective that these microstates arise from brain generators in these regions (see Supplementary Materials). Microstate C was associated with the medial temporal lobe subsystem, including sources in the precuneus, temporal lobes, and a large subcortical cluster extending to the hippocampus and parahippocampal cortex. Microstates D and E were associated with frontoparietal regions overlapping with dorsal attention and frontoparietal control networks (Dixon et al., 2018; Marek & Dosenbach, 2018; Spreng et al., 2013). Microstate D was associated with bilateral sources in inferior parietal lobule and dorsolateral prefrontal cortex, whereas microstate E was associated with sources in precuneus, motor and supplementary motor areas, and bilateral sources in middle frontal gyrus. Sources in middle frontal gyrus included the frontal eye fields and other regions implicated in attention and executive functions.

The maintenance of endogenous attention over the wait time delay of trials revealed substantial limitations in the span of individuals’ attention at shorter timescales. In both the current study and a prior behavioral study (Zanesco et al., 2026b), we found that the endogenous maintenance of attention was limited to just a few seconds before the likelihood of detection errors and mind wandering increased with time during the wait time delay. While microstate E was associated with correctly detecting the presentation of the target stimulus on trials, we also found that microstate E increased in its prevalence and occurrence rate over sequential epochs of the wait time delay. These findings implicate microstate E in the maintenance of endogenous attention; microstate E occurs more frequently on later epochs of the wait time delay. This suggests that as the demand imposed on the maintenance of attention increases over longer durations, the brain generators of microstate E activate more frequently. Microstates were also more likely to transition to microstate E in sequence as time increased over the wait time delay. Specifically, transitions of microstate B to other microstates were interrupted more frequently by the occurrence of microstate E.

One interpretation of these findings is that microstate E participates in the “short-cycle refresh system” (Robertson et al., 1997) proposed to maintain endogenous attention and support task performance by repeatedly reactivating task schemas in the absence of exogenous influences (Langner & Eickhoff, 2013). Stuss et al. (1995) suggests that task schemas require continual reactivation because they intrinsically decay over seconds and can be inhibited by competing task-irrelevant schemas. As attention is maintained over longer periods of time, brain systems supporting attention may require more frequent reactivation as demands accrue and shorten its refresh cycle. The occurrence of microstate E roughly twice a second is consistent with the short cycle of such a system, and increased occurrence rate of microstate E may indicate these brain systems have to work harder to maintain attention during longer wait time delays. Attention maintained over longer inter-target intervals has also been shown to increase activity in a network of frontal, parietal, and subcortical regions, including areas in the middle frontal gyrus, supplementary and pre-supplementary motor areas, anterior and posterior cingulate, cuneus and precuneus, insula, and thalamus (Breckel et al., 2011). These regions overlapped with those identified in a coordinate-based meta-analysis of vigilant attention (Langner & Eickhoff, 2013) and some of the intracranial sources of microstate E identified in the present study through source localization. Together, these patterns align with the general tendency for activity in regions of frontal cortex that support attention and executive functions to scale with task demand and difficulty until capacity is exceeded (Braver et al., 1997; Van Snellenberg et al., 2015).

The tendency for performance to worsen over a task session has come to characterize the consequential limitations in our ability to sustain attention over time. Increases over time in the occurrence of mind wandering parallel these decrements and correlate with intraindividual change in stimulus detection accuracy and response time variability in continuous performance tasks (Schwartzman et al., 2025; Thomson et al., 2014; Zanesco et al., 2024, 2025), suggesting common neurocognitive processes may be involved in increasing rates of stimulus detection errors and the occurrence of mind wandering. In line with these patterns, we found that reports of focus decreased and task-unrelated thought increased over the task session of this study and in a prior behavioral study employing the Sustained Attention to Cue Task (Zanesco et al., 2026b). Importantly, we found that microstate C also demonstrated within-task change over the course of the task session. Specifically, microstate C explained more GEV, occupied more of epochs’ time series, and had longer duration occurrences as the task progressed over trials. Microstate C was therefore less prevalent at the beginning of the task compared to the end. The demand imposed on sustained attention from prior task trials also appears to influence microstates in subsequent moments by increasing the incidence of microstate C on trials following those with long wait time delays. While microstate C was uniquely associated with change in microstate dynamics over the task session, all microstates also increased in global field power with time-on-task. This suggests at least two differentiable neurocognitive processes associated with waning sustained attention over the task session: increasing dominance of brain circuits associated with microstate C and global increases in amplitude of activity associated with all microstates.

Self-reports of focus were negatively associated with the prevalence and occurrence rate of microstate C, consistent with patterns differentiating incorrectly detected trials from correct trials and prior research comparing off-task to on-task reports (i.e., Zanesco et al., 2021). Associations with task-unrelated thought, however, were inconsistent with these broader patterns. Instead, we found that microstates D and E tended to occur more frequently and occupy more of the EEG time series when individuals reported experiencing more task-unrelated thought, while microstates A and F occurred less frequently and occupied less of the time series. While we expected microstates D and E to be associated with states of focus rather than mind wandering, one interpretation may be that these patterns are related to the recruitment of attention and cognitive control systems for the maintenance and manipulation of internally generated mental representations during mind wandering (Dixon et al., 2018; Fox et al., 2015; Spreng et al., 2010). This possibility aligns with a model of mind wandering and spontaneous thought emphasizing that task-unrelated thoughts are not merely the result of failures of attention but also reflect the cognitive processes by which dorsal attention and frontoparietal control networks work to maintain and guide internal streams of ongoing thought (Christoff et al., 2016).

Prestimulus power in the alpha frequency band is associated with states of inattention. Alpha power tends to be greater in the moments preceding stimulus detection errors in attention tasks (van Dijk et al., 2008; Michail et al., 2022; Samaha et al., 2020) or during episodes of mind wandering (Kam et al., 2022). One secondary goal of our study was to investigate whether the activity of microstates can account for these associations. We found that the spatial distribution of alpha power was dependent on the concurrent microstates at those same moments in time (see Supplementary Materials). This supports the notion that the brain generators of microstates are prominent generators of alpha power on the scalp. Yet it is also important to note that the generators of microstates contribute broadband power in other frequencies considered to be important for human cognition, such as the theta band. The prevalence of specific microstates in the EEG time series will therefore differentially contribute to estimates of average power at different electrode sites. Accordingly, differences in pre-stimulus alpha power reported in studies of attention likely result from the presence of distinct microstates. These considerations add to the growing recognition that traditional frequency-based approaches for the analysis of spontaneous EEG provide only a limited perspective on the millisecond dynamics of the brain (Donoghue et al., 2022; van Vugt 2007).

Additionally, we found that alpha power was lower on correct relative to incorrect trials at moments coincident with microstates and at electrode locations aligned with their spatial configuration. Alpha power increased globally over the course of the task session at most electrode locations, consistent with prior studies investigating change in alpha power with time-on-task (e.g., Kopčanová et al., 2025; Pershin et al., 2023), and across sequential epochs at the location of the absolute voltage maxima of microstate E. These differences are consistent with observed patterns in the global field power of microstates and the broader literature linking increased alpha power to inattention and mind wandering. When examining the specific microstates coincident with periods of sustained, large amplitude alpha oscillations, we found that microstate C was present to a greater degree on trials where individuals failed to detect the target, trials where the wait time was longer in the prior trial, and later trials of the task, suggesting the brain generators of microstate C oscillate more frequently in these contexts. Other microstates were also present during sustained alpha oscillatory episodes, more so on later trials than earlier in the task. These findings suggest that the amplitude and oscillatory behavior of the brain generators of microstates contribute substantial amounts of power in the alpha frequency band and are likely responsible for pre-stimulus differences in power in the moments preceding stimulus detection errors reported in prior studies.

Theories of the functional significance of alpha oscillations emphasize their role in modulating cortical excitability through inhibiting and controlling the timing of neural activity by rhythmically entraining firing to oscillations (Jensen & Mazaheri, 2010; Klimesch, 2012; Klimesch et al., 2007). The oscillatory regimes of microstates likely arise from the mechanisms by which the brain implements regionally synchronized coactivation or inhibition of neuronal populations. This enables the connectivity architectures that cycle from one microstate to another to coordinate or suppress activity dynamically throughout the brain in support of neurocomputation and ongoing cognition. As microstates demonstrate both periodic and aperiodic behavior, it is important that future studies directly investigate the functional significance of transient occurrences of microstates relative to instances when they persist in time by oscillating in polarity over the course of several oscillatory cycles. Furthermore, as the brain generators of microstates are prominent generators of alpha power, it is critical that existing theories of the functional significance of alpha oscillations begin to consider the growing evidence linking specific microstates and their dynamics to distinct aspects of cognition.

Our findings contribute to understanding how the millisecond dynamics of interacting brain electric microstates support the endogenous maintenance of focus and contribute to its inevitable lapses over time. In line with prior behavioral results (Zanesco et al., 2026b), our study provides additional neural evidence to suggest that mechanisms responsible for failures in the maintenance of attention at shorter timescales, such as when visual attention is momentarily maintained at a spatial location for just a few seconds or minutes, may be separable from those contributing to gradual degradation in task performance over longer timescales. While microstates offer a unique, complementary window into the rapid temporal dynamics of attention and the brain systems supporting these cognitive functions, understanding their functional significance will require the integration of multi-modal neuroimaging techniques to corroborate the brain generators of microstates and their links to large-scale functional brain networks. Source localization is inherently indeterminate; requiring that future studies examine patterns of brain activity in the SACT using neuroimaging to connect activity patterns to microstates and better understand the distributed brain regions involved in the maintenance of attention over short and long timescales.

## Acknowledgements

We thank Jason Tsukahara and the laboratory of Randall Engle at the Georgia Institute of Technology for making the Sustained Attention to Cue Task freely available for use as part of the toolbox of attention control tasks. We utilized the freely available Cartool EEG software toolbox (cartool.unige.ch) developed by Denis Brunet at the Functional Brain Mapping Lab, then at the Epilepsy and Networks Lab, University of Geneva, Switzerland, and supported by the Center for Biomedical Imaging (CIBM), Switzerland.

## Data Availability Statement

We report all data exclusions, manipulations, and measures included in this study. The study was not preregistered. The data that support the findings of this study are available in the OSF repository and can be found at: https://osf.io/pw3e8

## Supplementary Materials

### Microstate Averaged Time-Frequency Transformation

Time-frequency transformation (“wavelet” S-Transform implemented in Cartool; Brown et al., 2010) of the spatially smoothed EEG time series of epochs determined the power in the alpha (8–14 Hz) frequency bands for all electrodes. This enabled estimation of localized variations of alpha power in time within each time series of epochs. Power in this band was subsequently averaged for each epoch according to the microstate configurations categorizing the time samples of the EEG. Power at uncategorized periods of time was ignored. The result was an estimate of power for all electrodes and microstate configurations for each of the 14,479 epochs. Supplementary Figure 1 depicts an example of the results of the S-transform of the continuous voltage waveforms, providing a representation of alpha power over time for all electrode locations for a single 4000 msec epoch. The topographic distribution of power is shown averaged according to the categorized time series of microstates.

Episodes of sustained, large amplitude alpha oscillations were also identified using an approach motivated by methods for identifying oscillatory episodes from wavelet transformed EEG (Hughes et al., 2012; van Vugt et al., 2007). Across all epochs for an individual, time samples of alpha power (averaged across all electrodes) with amplitude greater than the 70th percentile were identified. Time samples in the 70th percentile had to be sustained over time for several oscillatory cycles. Episodes of large amplitude alpha were therefore only identified as an oscillatory episode if they were sustained in time for at least 200 msec. The number of time samples that distinct microstates were present within these episodes was quantified and compared between microstates. Supplementary Figure 1 depicts the identified alpha oscillatory episodes present for a single 4000 msec epoch and their overlap with the categorized time series of microstates.

**Supplementary Figure 1.**
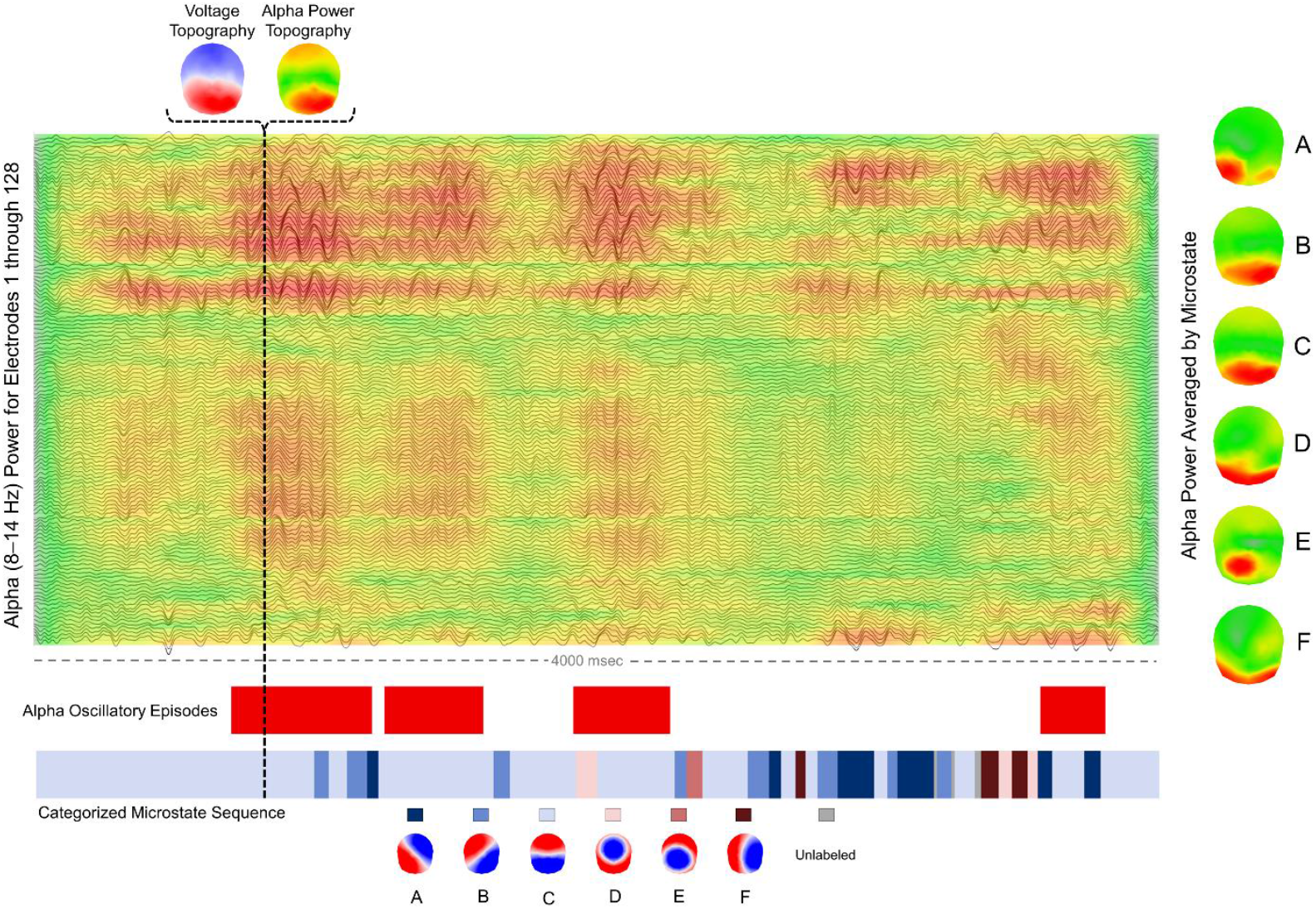
Alpha (8–14 Hz) power is shown for the 128-channel EEG of a single 4000 msec epoch during the wait time delay of a trial. The voltage waveforms for all 128 channels are also shown aligned to power estimates. The categorized microstate time series is provided below the epoch, indicating each time series sample categorized as a particular microstate configuration. The voltage topography and alpha power topography are shown for a single example time sample. The sample was categorized as microstate configuration C and both topographies resemble configuration C. Alpha power is averaged in time according to the categorized time series of microstates to generate maps of power corresponding to each microstate for every epoch of task trials. The spatial distribution of average alpha power corresponding to each microstate is shown on the right for this epoch.

### Analysis

Microstate averaged alpha and theta power. Power in the alpha (8–14 Hz) and theta (4–8 Hz) bands were averaged at each electrode location according to the microstates categorizing time samples of the EEG for each epoch. We compared averaged alpha and theta power at each electrode location between microstate configurations to examine whether the spatial distribution of power was dependent on microstate configuration. Next, we examined average alpha and theta power as a function of trial accuracy, change over epochs of the wait time delay, and change over sequential trials of the task. Mixed effects models were estimated with restricted maximum likelihood using the PROC HPMIXED procedure in SAS 9.4. We set the significance level of all model parameters in these mixed effects models to α = 0.001.

## Supplementary Results

### Distributed Brain Sources of Microstates

Source localization revealed distributed networks of brain regions contributing to the scalp electric field configuration of microstates. Clusters of sources above the 95th percentile were in regions that overlapped with key areas of large-scale brain networks with functional significance for sustained attention and mind wandering. Average patterns of sources are summarized in Supplementary Figure 2 for each microstate configuration. More comprehensive depictions of neural sources are depicted across a range of sagittal, axial, and coronal slices in Supplementary Figures 3 through 5.

**Supplementary Figure 2.**
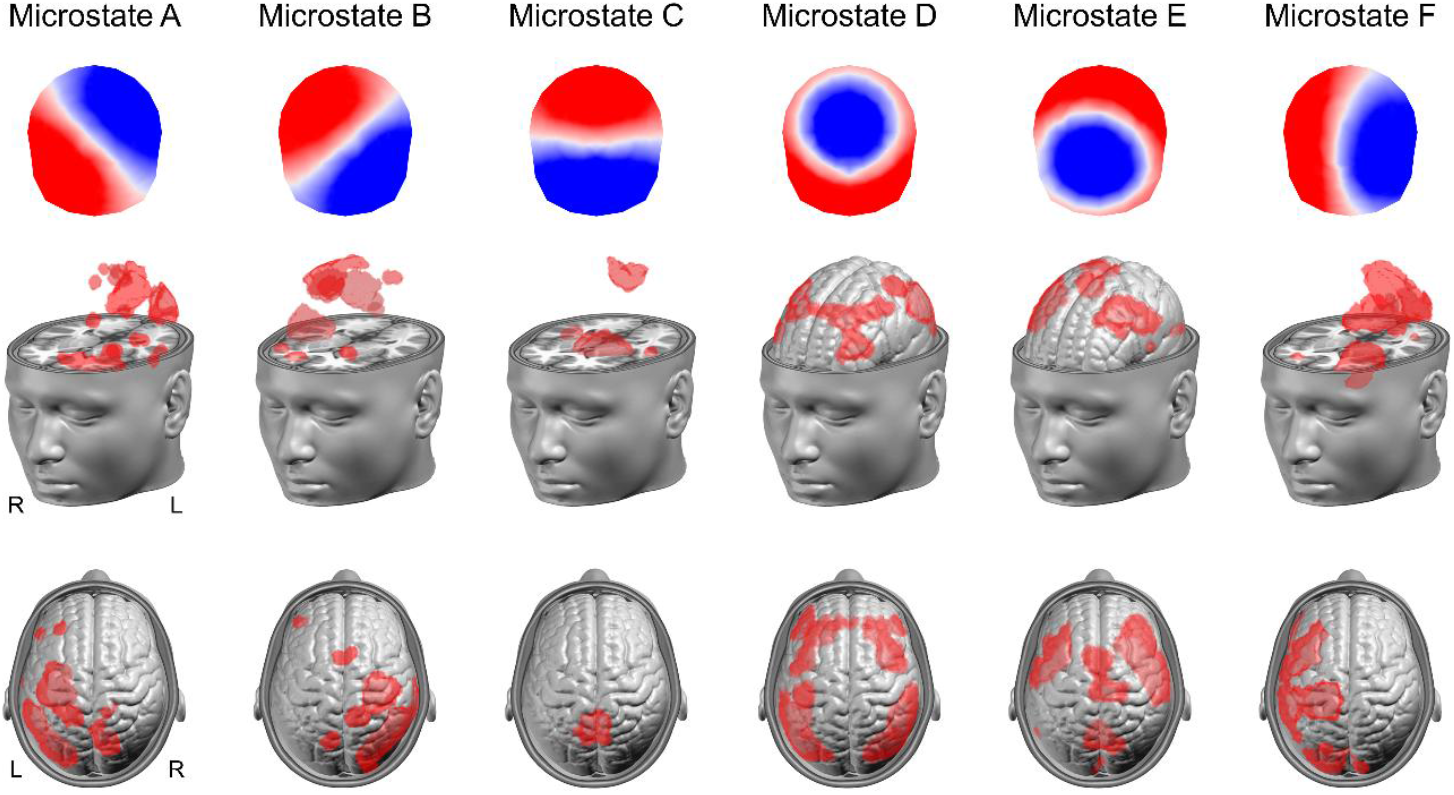
Neural sources of microstates are summarized from distributed source localization of EEG. Sources are thresholded to values above the 95th percentile and projected onto the MNI template brain. Microstate configurations are shown above each corresponding map of sources.

**Supplementary Figure 3.**
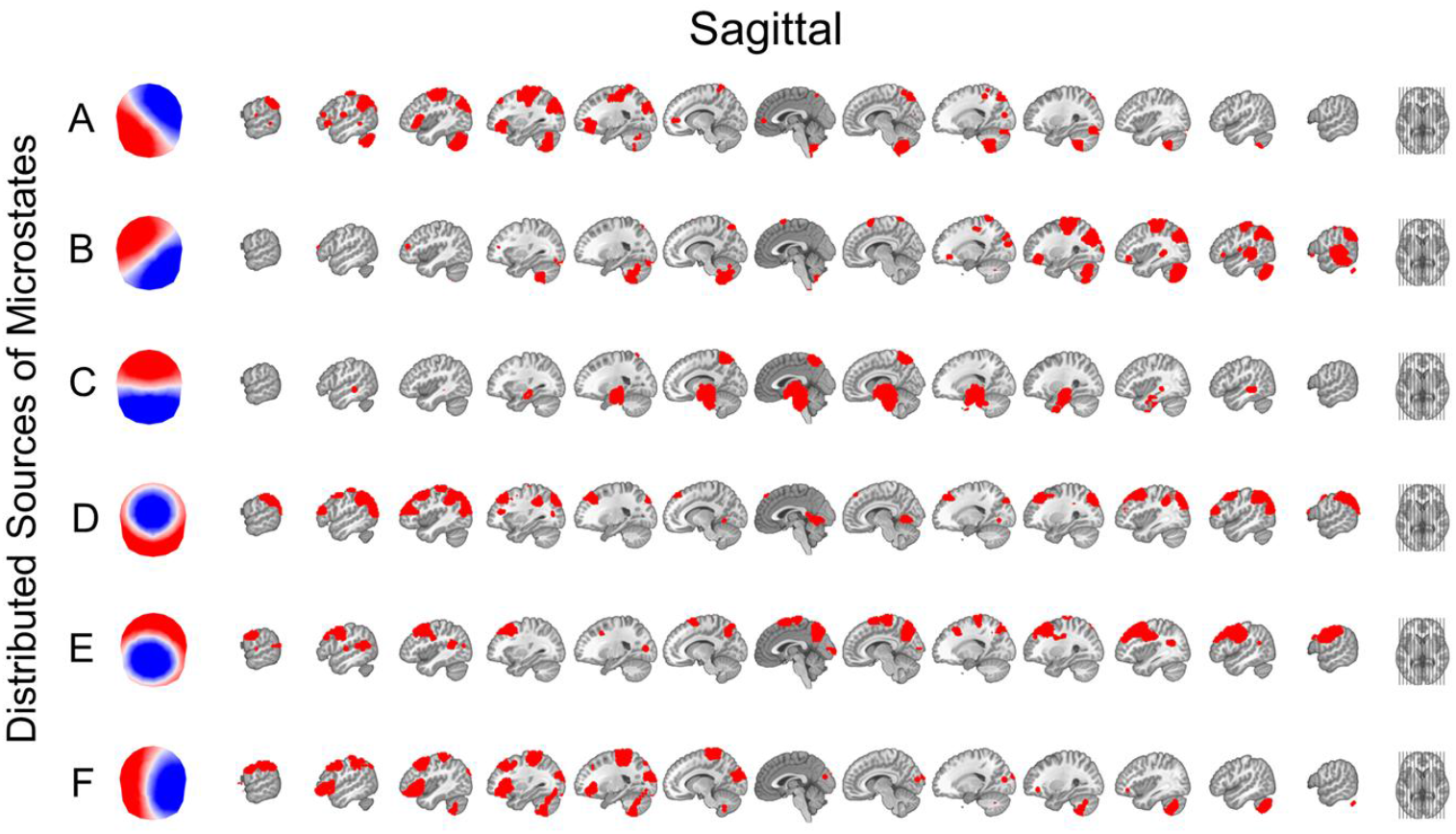
Distributed neural sources of microstates are summarized from source localization of EEG. Sources are thresholded to values above the 95th percentile and projected onto the MNI template brain for a range of sagittal slices.

Microstate A was associated with sources in the cerebellum, cuneus, left inferior occipital gyrus, left superior occipital gyrus and precuneus, left inferior parietal lobule, left pre and postcentral gyri, and left middle and inferior frontal gyri. Additionally, there were smaller clusters of sources in medial frontal gyrus, right precuneus, left middle frontal gyrus, and left middle and superior temporal gyri. Microstate B was associated with sources in cerebellum, right cuneus and middle occipital gyrus, right precuneus and inferior parietal lobule, right pre and postcentral gyri, right superior, middle, and inferior temporal gyri, right middle and inferior frontal gyri. Smaller clusters were also in superior frontal gyrus, left superior parietal lobule, and left middle frontal gyrus. Microstate C was associated with sources in precuneus, right and left superior and middle temporal gyrus, and an extensive cluster of subcortical sources extending to the medial temporal lobes. Microstate D was associated with bilateral sources in the inferior parietal lobules and bilateral sources in pre and postcentral gyrus, middle frontal gyrus, and superior frontal gyrus. Finally, there was a small cluster of sources in the cuneus and lingual gyrus. Microstate E was associated with sources in precuneus, precentral and superior frontal gyrus, and bilateral sources in middle frontal gyrus and superior frontal gyrus. Smaller clusters were found in cuneus and left and right superior temporal gyrus. Microstate F was associated with sources in cerebellum and cuneus, left pre and postcentral gyrus, left inferior parietal lobule, left middle frontal gyrus, and a large cluster in the left inferior frontal gyrus.

### Microstate Averaged Alpha Power

Power in the alpha (8–14 Hz) band was averaged at each electrode location according to the microstates categorizing time samples of the EEG. We first compared averaged alpha power at each electrode location between microstate configurations to examine whether the spatial distribution of power was dependent on microstate configuration. Supplementary Figure 6 (left) depicts the microstate averaged power maps for each configuration alongside maps of significant (p < .001) t-values from comparisons between configurations. Importantly, power maps recapitulated the voltage map of each microstate configuration. The spatial distribution of power was therefore explained by concurrent microstates. Time series samples labeled as configuration C contributed the most alpha power to anterior and posterior scalp electrode locations. Configuration A and B contributed additional power to left and right posterior electrode locations, respectively. Power was greatest for samples coincident with configuration D at fronto-central locations and greatest for configuration E at posterior-central locations.

Alpha power at each electrode location was next examined as a function of trial accuracy, change over epochs of the wait time delay, and change over sequential trials of the task. Supplementary Figure 6 (right) depicts the microstate averaged power maps for correct and incorrect trials alongside maps of significant (p < .001) t-values for simple effects of trial accuracy, epoch, and sequential trial number for all microstate configurations. Alpha power was lower for correct vs. incorrect trials at many electrode locations for all microstate configurations. The spatial distribution of significant electrodes aligned with the spatial configuration of microstates. Microstate configuration C demonstrated the largest differences in power across anterior and posterior electrode sites. Alpha power was greater across sequential epochs at only a few electrode locations. The distribution of alpha power effects across sequential epochs was notably located at the topographic location where the absolute value of the voltage map of microstate E was maximal (i.e., the antinode of the standing wave), suggesting these effects may reflect increases in alpha power resulting from specific oscillations in the polarity of microstate E. Finally, alpha power increased over sequential task trials at many electrode locations for all microstate configurations. These effects were also maximal at electrode locations coincident with the spatial configuration of microstates.

**Supplementary Figure 4.**
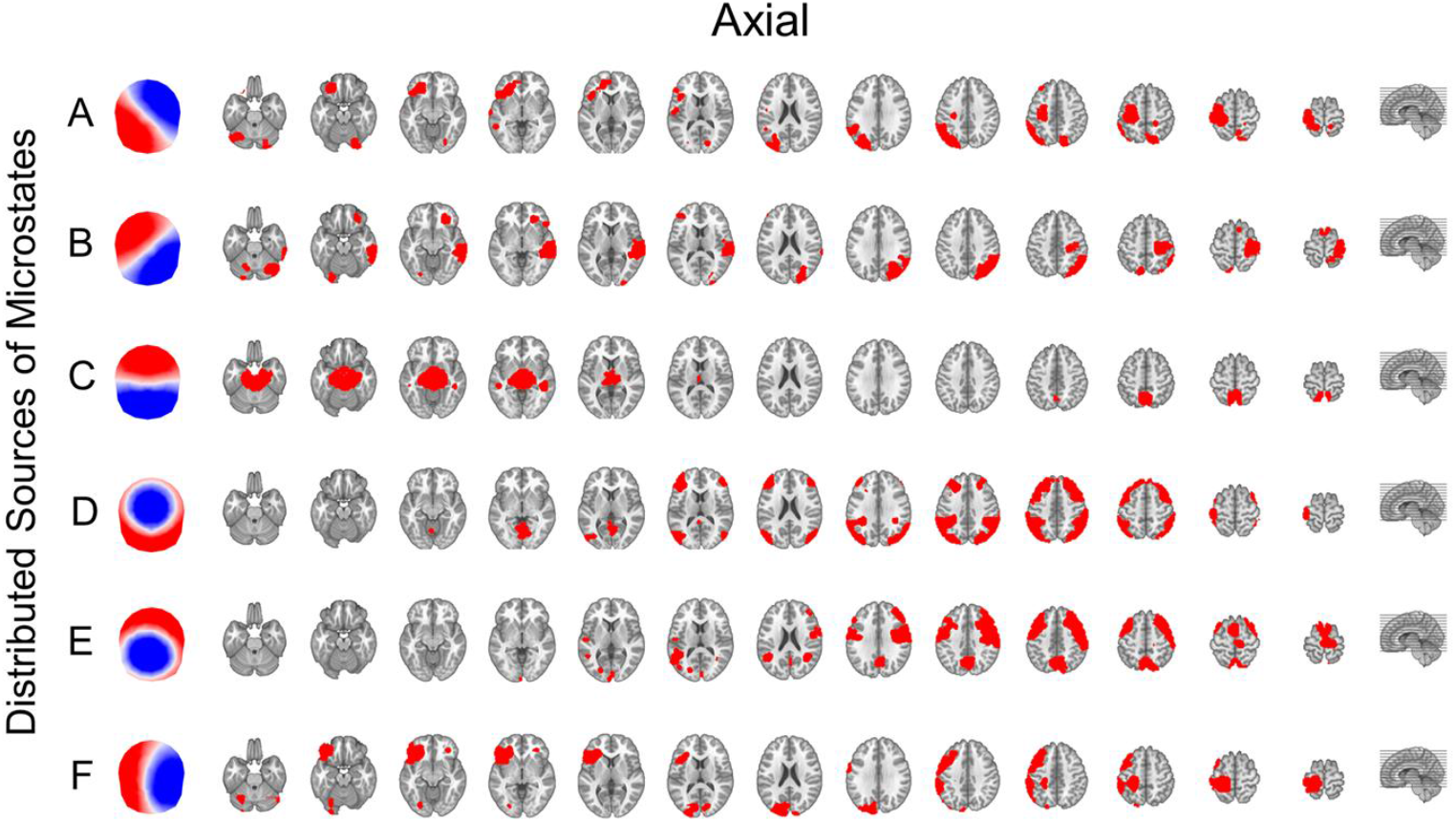
Distributed neural sources of microstates are summarized from source localization of EEG. Sources are thresholded to values above the 95th percentile and projected onto the MNI template brain for a range of axial slices.

**Supplementary Figure 5.**
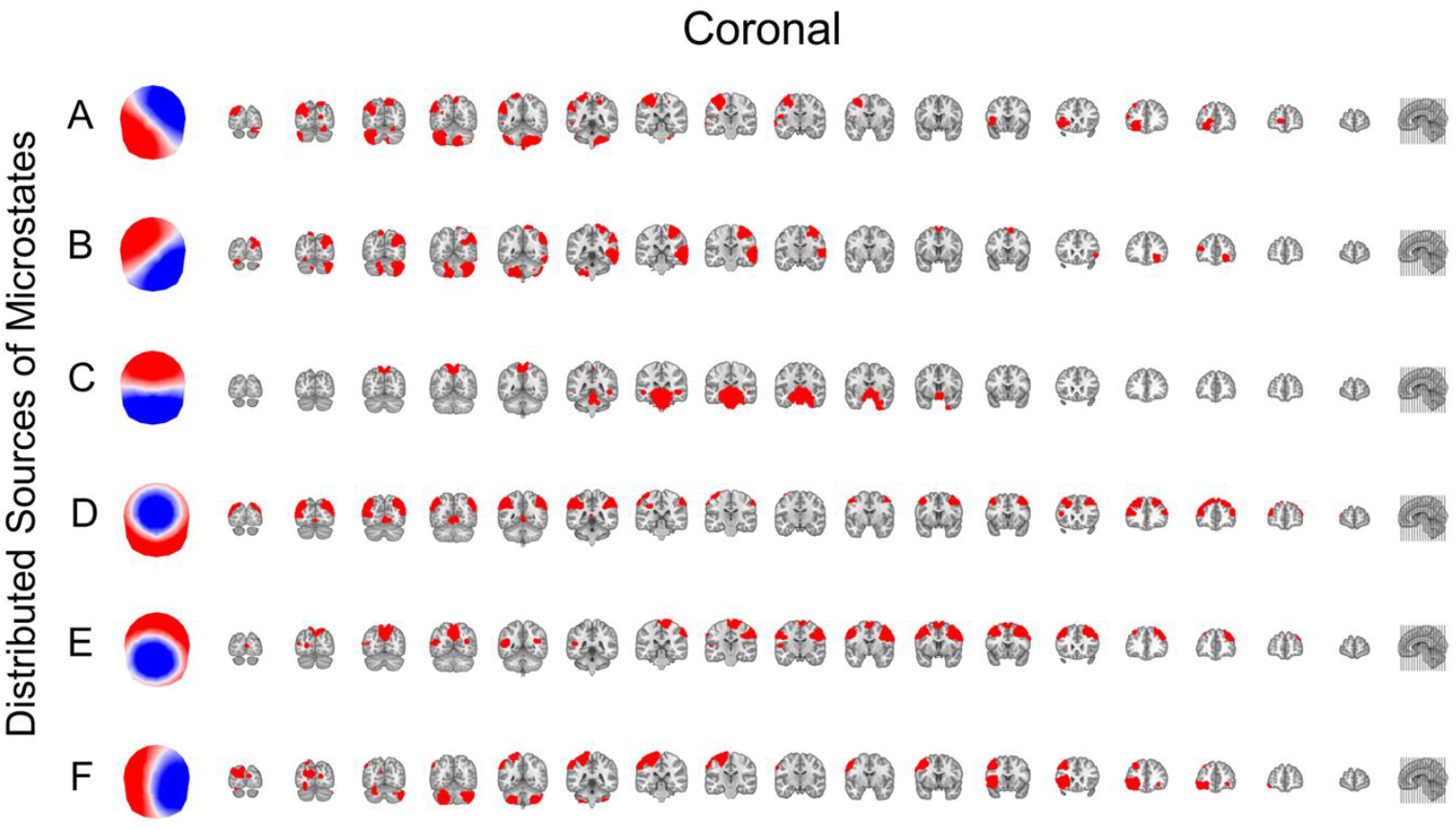
Distributed neural sources of microstates are summarized from source localization of EEG. Sources are thresholded to values above the 95th percentile and projected onto the MNI template brain for a range of coronal slices.

**Supplementary Figure 6.**
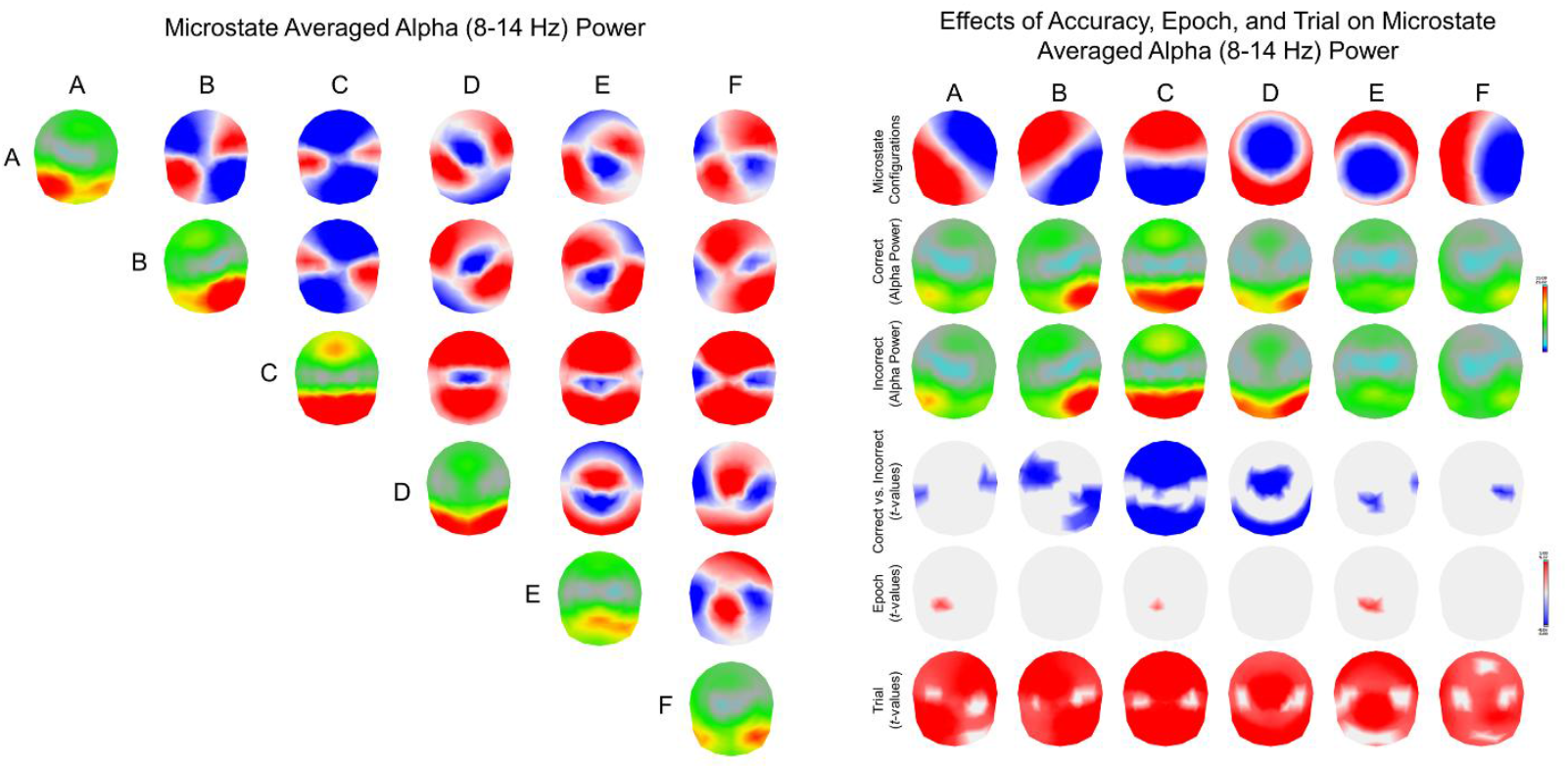
The spatial distribution of average alpha power (8–14 Hz) for each microstate configuration is depicted from mixed-effects model estimates comparing microstate configurations on the left, and the effects of accuracy, sequential epoch, and sequential trial on the right. On the left, model estimates of mean alpha power are plotted for all electrode locations on the diagonal for each microstate configuration (red and yellow is more power at that location). Maps on the upper diagonal provide significant (*p* < .001) *t*-values for electrode locations from pairwise comparisons between configurations on rows compared to configurations in columns. On the right, maps display mean power estimates partitioned by correct and incorrect trials, separately for each microstate configuration. Maps of significant (*p* < .001) *t*-values illustrate electrodes with differences between correct and incorrect trials, significant change over epochs of the wait time delay, and significant change over sequential trials of the task. Across all comparison maps, positive *t*-values are shown in red (indicating greater power at that electrode location), while negative values are shown in blue. Non-significant *t*-values (*ps* > .001) are omitted.

#### Alpha oscillatory episodes

We identified the number of time samples that microstates were present within windows of sustained, large amplitude alpha oscillations using a modified oscillatory episode detection method (cf. Hughes et al., 2012; van Vugt et al., 2007). We examined whether the number of time samples within oscillatory episodes significantly differed between microstates by evaluating the effects of microstate configuration across all epochs in a mixed-effects model. There was a significant effect of microstate configuration, F(5, 57303) = 3508.62, p < .001, indicating that some microstates were more present within oscillatory episodes compared to others. On average across epochs, microstate configuration A was present 44.3772 (SE = 1.8644, 95% CI [40.7230, 48.0314]) time samples, whereas microstate B was present 61.0583 (SE = 1.8485, 95% CI [57.4352, 64.6814]) time samples, microstate C was present 132.2331 (SE = 1.8353, 95% CI [128.64, 135.83]) time samples, microstate D was present 52.1413 (SE = 1.8533, 95% CI [48.5088, 55.7738]) time samples, microstate E was present 53.6132 (SE = 1.8634, 95% CI [49.9610, 57.2654]) time samples, and microstate F was present 32.4631 (SE = 1.8936, 95% CI [28.7517, 36.1746]) time samples. As such, microstate C was present within oscillatory episodes significantly more than all other microstates (all ps < .001), occupying more than twice as many time samples within windows. These findings suggest that the brain generators of microstate C were also the most prominent generators of sustained alpha oscillations in the EEG.

We next examined the effects of trial accuracy, sequential epoch, sequential trial number (i.e., time-on-task), and their two-way interactions with microstate configuration by including these effects in a second mixed-effects model. We observed a significant main effect of microstate configuration, F(5, 43348) = 261.92, p < .001, no significant main effect of trial accuracy, F(1, 43348) = 3.61, p = .057, no significant main effect of epoch, F(1, 43348) = 0.37, p = .543, and a significant main effect of sequential trial number, F(1, 43348) = 29.17, p < .001. Importantly, we also observed a significant interaction between trial accuracy and microstate configuration, F(5, 43348) = 3.32, p = .005, and a significant interaction between trial number and microstate configuration, F(5, 43348) = 47.99, p < .001. There was no significant interaction between sequential epoch and microstate configuration, F(5, 43348) = 2.48, p = .029.

We found that the time occupied by microstate C in oscillatory episodes differentiated correctly detected trials from incorrect trials. Microstate configuration C occupied significantly fewer time samples within oscillatory episodes (b = −5.9387, SE = 1.3337, p < .001, 95% CI [−8.5528, −3.3247]) on epochs of correct trials on average relative to incorrect trials, suggesting that the generators of microstate C oscillated less frequently on correct trials. Several microstates also occupied more time samples within oscillatory episodes as the task progressed in time during the session. Microstate configuration B (b = 0.3084, SE = 0.0721, p < .001, 95% CI [0.1671, 0.4496]), microstate configuration C (b = 0.8653, SE = 0.0704, p < .001, 95% CI [0.7272, 1.0033]), microstate configuration D (b = 0.3553, SE = 0.0727, p < .001, 95% CI [0.2129, 0.4978]), and microstate configuration E (b = 0.2545, SE = 0.0737, p < .001, 95% CI [0.1100, 0.3991]), all occupied more time samples within oscillatory episodes of epochs as the task progressed in time. These microstates were therefore present within windows of sustained, large amplitude alpha oscillations to a greater degree on incorrect trials and later trials of the task, suggesting the generators of microstates oscillated more frequently. Finally, including the effects of previous trial wait time in a third mixed-effects model revealed a significant interaction between previous trial wait time and microstate configuration, F(5, 42828) = 3.02, p = .010. Microstate configuration C occupied more time samples within oscillatory episodes (b = 0.2531, SE = 0.0544, p < .001, 95% CI [0.1465, 0.3596]) of current trial epochs when the wait time was longer in the previous trial.

### Microstate Averaged Theta Power

Power in the theta (4–8 Hz) band was averaged at each electrode location according to the microstates categorizing time samples of the EEG. We first compared averaged theta power at each electrode location between microstate configurations to examine whether the spatial distribution of power was dependent on microstate configuration. Supplementary Figure 7 (left) depicts the microstate averaged power maps for each configuration alongside maps of significant (p < .001) t-values from comparisons between configurations. Importantly, power maps recapitulated the voltage map of each microstate configuration. The spatial distribution of power was therefore explained by concurrent microstates. Time series samples labeled as configuration C contributed the most theta power to anterior and posterior scalp electrode locations. Configuration A and B contributed additional power to left and right posterior electrode locations, respectively. Power was greatest for samples coincident with configuration D at fronto-central locations and greatest for configuration E at posterior-central locations.

Theta power at each electrode location was next examined as a function of trial accuracy, change over epochs of the wait time delay, and change over sequential trials of the task. Supplementary Figure 7 (right) depicts the microstate averaged power maps for correct and incorrect trials alongside maps of significant (p < .001) t-values for simple effects of trial accuracy, epoch, and sequential trial number for all microstate configurations. Theta power was lower for correct relative to incorrect trials at electrode locations coincident with the absolute voltage maxima of microstate configurations. Theta power was greater across sequential epochs at central electrode locations for all microstates. Finally, theta power increased over sequential task trials for all microstate configurations at electrode locations coincident with the spatial configuration of microstates.

**Supplementary Figure 7.**
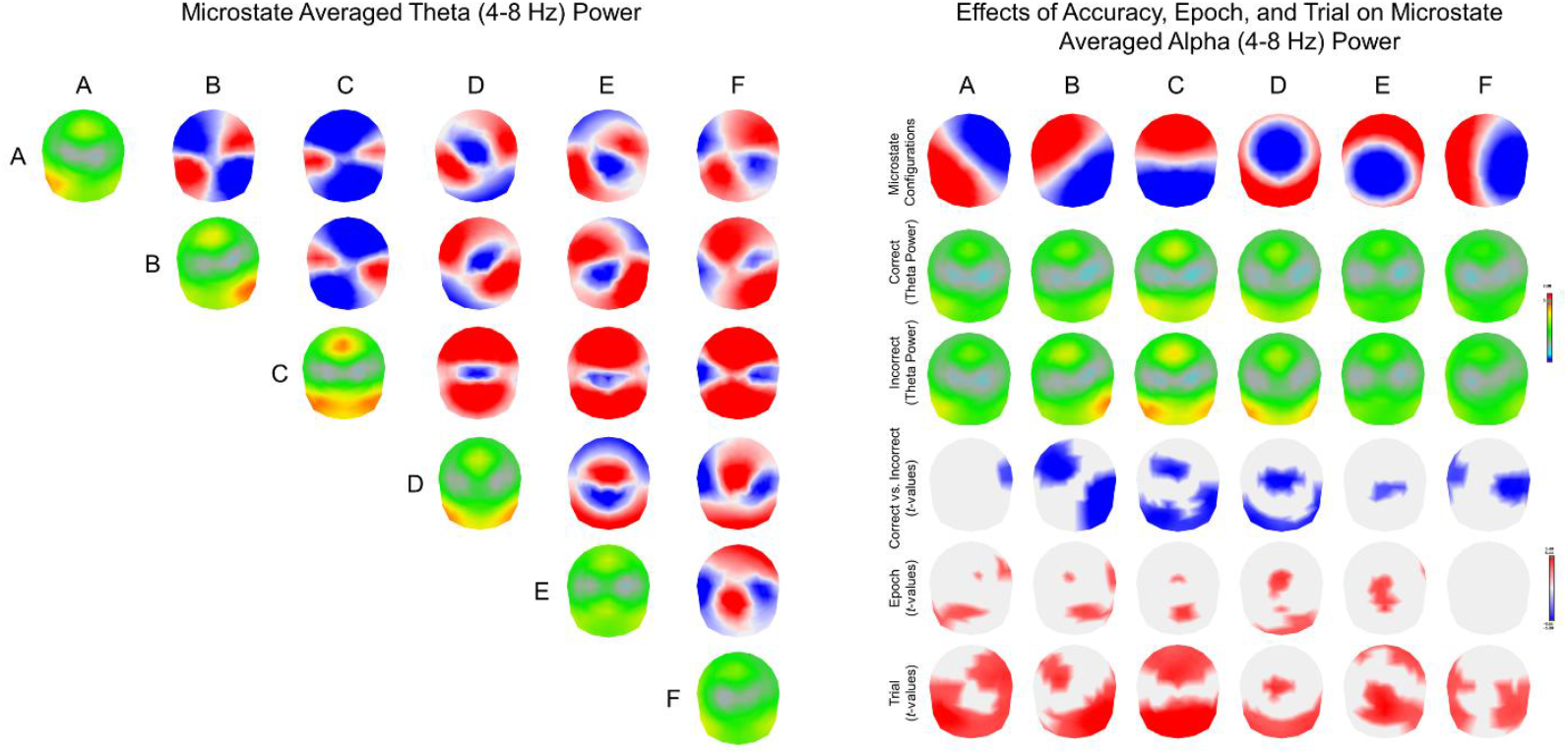
The spatial distribution of average theta power (4–8 Hz) for each microstate configuration is depicted from mixed-effects model estimates comparing microstate configurations on the left, and the effects of accuracy, sequential epoch, and sequential trial on the right. On the left, model estimates of mean theta power are plotted for all electrode locations on the diagonal for each microstate configuration (red and yellow is more power at that location). Maps on the upper diagonal provide significant (*p* < .001) *t*-values for electrode locations from pairwise comparisons between configurations on rows compared to configurations in columns. On the right, maps display mean power estimates partitioned by correct and incorrect trials, separately for each microstate configuration. Maps of significant (*p* < .001) *t*-values illustrate electrodes with differences between correct and incorrect trials, significant change over epochs of the wait time delay, and significant change over sequential trials of the task. Across all comparison maps, positive *t*-values are shown in red (indicating greater power at that electrode location), while negative values are shown in blue. Non-significant *t*-values (*ps* > .001) are omitted.

